# Lysosomal retargeting of Myoferlin mitigates membrane stress to enable pancreatic cancer growth

**DOI:** 10.1101/2021.01.04.425106

**Authors:** Suprit Gupta, Julian Yano, Htet Htwe Htwe, Hijai R. Shin, Zeynep Cakir, Thomas Ituarte, Kwun W. Wen, Grace E. Kim, Roberto Zoncu, David W. Dawson, Rushika M. Perera

## Abstract

Lysosomes must maintain integrity of their limiting membrane to ensure efficient fusion with incoming organelles and degradation of substrates within their lumen. Pancreatic cancer cells upregulate lysosomal biogenesis to enhance nutrient recycling and stress resistance, but whether dedicated programs for maintaining lysosomal membrane integrity facilitate pancreatic cancer growth is unknown. Using proteomic-based organelle profiling, we identify the Ferlin family plasma membrane repair factor, Myoferlin, as selectively and highly enriched on the membrane of pancreatic cancer lysosomes. Mechanistically, lysosome localization of Myoferlin is necessary and sufficient for maintenance of lysosome health and provides an early-acting protective system against membrane damage that is independent from the endosomal sorting complex required for transport (ESCRT)-mediated repair network. Myoferlin is upregulated in human pancreatic cancer, predicts poor survival, and its ablation severely impairs lysosome function and tumour growth *in vivo*. Thus, retargeting of plasma membrane repair factors enhances pro-oncogenic activities of the lysosome.

---

Lysosomes function as critical nodes for macromolecular recycling, vesicle trafficking, metabolic reprogramming, and pro-growth signaling in the cell^1–3^. Pancreatic ductal adenocarcinoma (PDA), are highly reliant on enhanced lysosome function to facilitate degradation, clearance and recycling of cellular material delivered by increased rates of vesicle trafficking through autophagy and macropinocytosis^4–9^. To cope with the increased flux of substrates delivered to the lysosome for degradation, PDA cells upregulate transcriptional programs for lysosome biogenesis, which are mediated by the MiT/TFE factors^6, 10^. Although enhanced MiT/TFE activity leads to a dramatic increase in the number of lysosomes^6^, whether qualitative differences endow PDA lysosomes with unique structural and functional properties to cope with higher rates of vesicle trafficking and substrate clearance remains unknown.

An emerging aspect is the susceptibility of lysosomes to undergo membrane damage due to various cellular stressors. In response to damage, several mechanisms ensure proper lysosomal integrity and function by responding to damage and facilitating repair or removal of dysfunctional lysosomes^11, 12^. These pathways include repair via the Endosomal Sorting Complex Required for Transport (ESCRT), a multi-subunit membrane-remodeling machinery that performs membrane bending and scission away from the cytoplasm^12^. ESCRT components were recently shown to polymerize on the limiting membrane of lysosomes challenged with membrane-damaging agents, where they facilitate repair of ‘microtears’ likely through scission and resealing of the free membrane edges^13, 14^. When ESCRT recruitment to damage sites is blocked, more permanent and severe damage to lysosome function results. A second pathway facilitates clearance of irreversibly damaged or dysfunctional lysosomes via a selective macroautophagy process, termed lysosphagy^11, 15–17^.

PDA cells are unusual in their ability to import and degrade a large variety and quantity of intracellular and extracellular substrates to the lysosomal lumen in order to sustain unrestrained growth in nutrient-poor microenvironments. While it is likely that this enhanced substrate intake may pose unique challenges, it remains unknown whether cancer-specific mechanisms for lysosomal membrane stabilization enable PDA cells to cope with higher susceptibility to organelle stress. To answer this question, we conducted mass spectrometry-based profiling of immuno-isolated lysosomes captured from PDA and non-PDA cells. We uncovered profound differences in the protein content and composition of PDA lysosomes and identified members of the Ferlin family of membrane repair factors, Myoferlin (MYOF) and Dysferlin (DYSF) as membrane proteins uniquely enriched in PDA lysosome fractions, with MYOF being broadly upregulated in PDA cell lines and patient samples. We find that lysosomal MYOF protects against a variety of membrane stressors to sustain enhanced functionality of PDA lysosomes. Accordingly, MYOF depletion in PDA cells triggers constitutive lysosomal membrane damage, leading to profound defects in lysosome morphology and function and arrested PDA tumour growth. Importantly, MYOF is upregulated in patient PDA specimens and high levels are predictive of poor patient prognosis, thus implicating MYOF as a key regulator of lysosome function and PDA tumour growth.

## Results

### Organelle proteomics identifies the Ferlin repair factors as PDA specific lysosomal membrane proteins

To identify novel proteins specific to PDA lysosomes, we captured intact lysosomes via affinity purification from cells stably expressing the lysosome membrane protein TMEM192 fused to mRFP and 3x HA tag (T192-mRFP-3xHA; LysoTag) (Fig. 1a; Supplementary Fig. 1a). Expression of LysoTag allows for efficient and rapid capture of intact lysosomes from cells that are amenable to mass spectrometry-based proteomics analysis^18, 19^. Comparative analysis of proteins present in lysosome fractions from PDA cells (PaTu8988T) versus non-PDA cells (HEK293T) identified 376 proteins with *≥* 2-fold enrichment in PDA lysosomal elutes (Supplementary Table 1). Analysis of the biological processes and pathways associated with these enriched PDA lysosomal proteins identified pathways associated with metabolism (“fatty acid degradation”, “valine, leucine, isoleucine degradation”, “central carbon metabolism in cancer”), as well as cell adhesion (“focal adhesion”, “ECM-receptor interaction”) (Fig. 1b). Moreover, we noted that proteins associated with “vesicle-mediated trafficking” (eg. RAB22A, SNX11) and “endocytosis” (eg. CLTB, DNML1, PACSIN3, ITSN1, EZR) were significantly enriched in PDA lysosome elutes consistent with heightened rates of vesicle trafficking converging on the lysosome in PDA cells (Fig. 1b,c; Supplementary Table 2). Moreover, several autophagy related proteins were enriched in PDA lysosomes, such as LC3B, GABARAP2 and autophagy receptors (NBR1, WDFY3, SEC62, TEX264, P62), consistent with increased rates of autophagy in PDA (Fig. 1d,e). As a control, levels of LAMP1 were similar in HEK293T and PaTu8988T lysosomes (Fig. 1d,e).

**Fig. 1.**
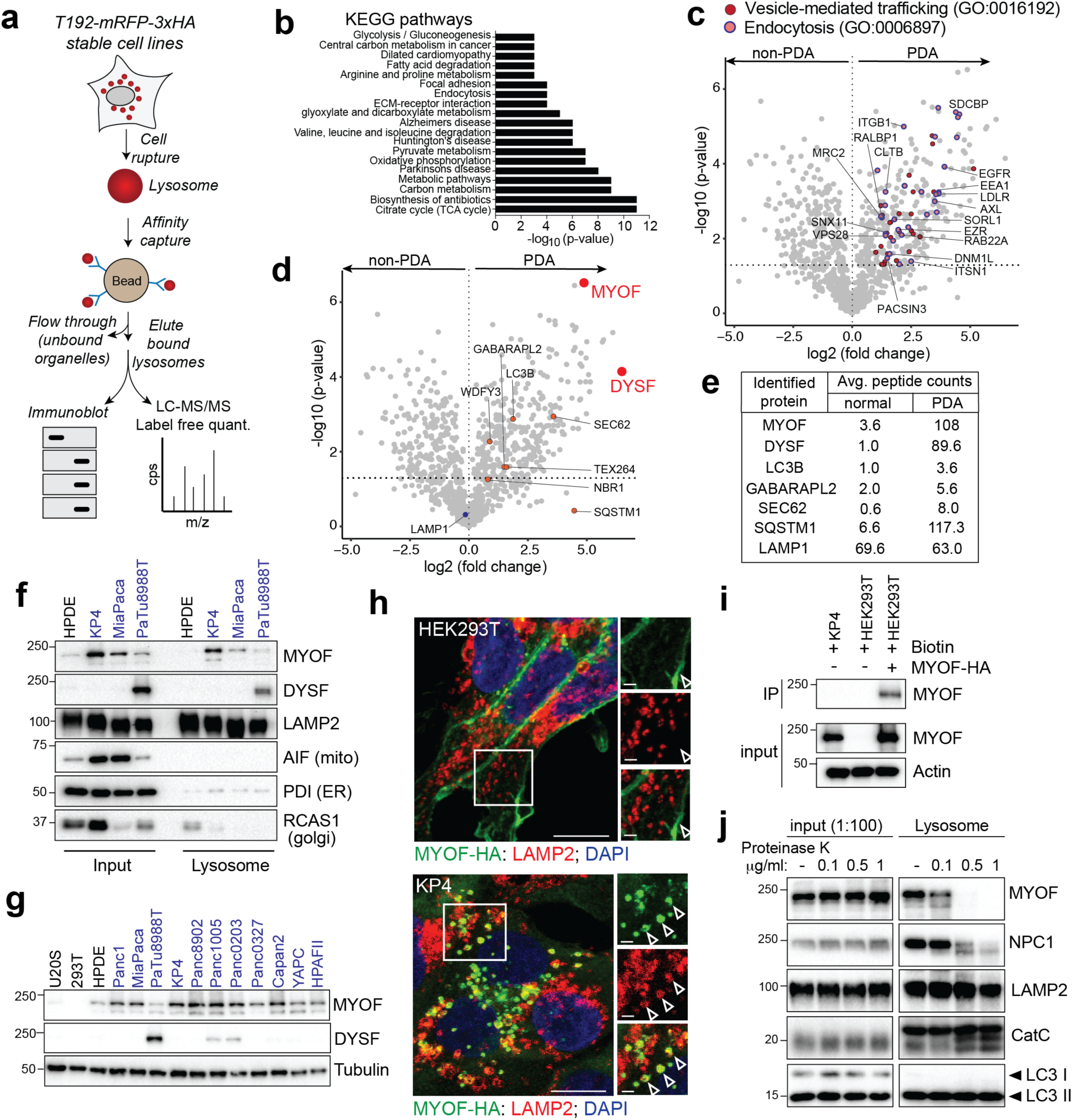
Organelle proteomics identifies the Ferlin repair factors as PDA specific lysosome associated membrane proteins. **a.** Schematic showing lysosome purification using affinity-based capture from cells stably expressing T192-mRFP-3xHA. **b.** KEGG pathway analysis of *≥*2-fold enriched PDA lysosome associated proteins. **c.** Volcano plot of lysosome proteomics data from non-PDA (HEK293T) and PDA (PaTu8988T) cells. Data are plotted as log2 fold change (PDA/non-PDA) versus the −log10 of the p-value. *≥*2-fold enriched proteins associated with “vesicle mediated trafficking” are indicated in dark red and overlapping proteins associated with “endocytosis” are indicated in pink/blue. **d.** Identical volcano plot as in (c) indicating autophagy related proteins (orange) and MYOF and DYSF (red). **e.** Average peptide counts for the indicated proteins from n = 3 biological replicates. **f.** Immunoprecipitation of purified lysosomes from the indicated cell lines showing enrichment of MYOF and DYSF in PDA lysosome fractions. LAMP2 serves as a loading control while absence of AIF, PDI and RCAS1 confirm organelle purity. **g.** Immunoblot showing levels of MYOF and restricted expression of DYSF in the indicated human cell lines (PDA highlighted in blue). **h.** Immuno-fluorescence staining of MYOF-HA (green) and LAMP2 (red) in HEK293T (left) and KP4 (right) cells. Arrowheads indicate plasma membrane localization of MYOF in HEK293T cells and lysosome localization in KP4 cells. Scale, 20μm, inset scale, 2μm. **i.** Biotinylation of cell surface proteins in KP4 and HEK293T cells expressing MYOF-HA. Biotinylated proteins were immuno-precipitated and western-blotted for MYOF. Note, MYOF is not on the cell surface of KP4 cells while MYOF-HA is present on the cell surface when expressed in HEK293T cells. **j.** Affinity purified lysosomes were treated with increasing concentrations of Proteinase K as indicated. Intraluminal proteins are protected from degradation (LAMP2, Cathepsin C, LC3B) while extra-luminal proteins are sensitive to digestion (NPC1 and MYOF).

Interestingly, two of the most significantly enriched proteins in PDA lysosome fractions were Myoferlin (MYOF; 30-fold enriched) and Dysferlin (DYSF; 89.6-fold enriched) (Fig. 1d,e), both of which belong to the Ferlin family of membrane repair factors^20–22^. We first validated the PDA-specific enrichment of MYOF and DYSF via direct immunoblotting of lysosomal fractions from HPDE (human pancreatic ductal epithelial cells) and a panel of PDA cells expressing LysoTag. Consistent with our proteomics data, PaTu8988T lysosome fractions expressed both MYOF and DYSF (Fig. 1f). Lysosomes isolated from KP4 and MiaPaca cells also displayed high levels of MYOF relative to HPDE lysosomes (Fig. 1f). Moreover, lysosomal enrichment of Ferlins correlated with their elevated expression in multiple PDA lines, whereas these proteins showed low or undetectable expression in a panel of non-PDA cells (Fig. 1g). Of note, MYOF transcript and protein levels were broadly upregulated in virtually all human PDA cell lines while DYSF upregulation was restricted to 3 cell lines (**Supplementary** figure 1b) with PaTu8988T cells having higher levels of DYSF relative to MYOF (Fig. 1g).

Prior studies have shown that Ferlin proteins localize predominantly to the plasma membrane, where they facilitate repair of the lipid bilayer in tissues subjected to heightened mechanical stress, particularly in skeletal muscle cells^21, 23, 24^. Accordingly, mutations in DYSF are associated with two forms of muscular dystrophy; Limb Girdle Muscular Dystrophy type 2B (LGMD2B) and Miyoshi Myopathy (MM), whereby impaired membrane resealing compromises myoblast maturation, fusion and plasma membrane repair^20, 25, 26^. Our lysosomal mass spectrometry analysis suggests that, in PDA cells, Ferlin proteins may be retargeted to the lysosomal membrane. Consistent with this hypothesis, transient expression of MYOF-HA in PDA cells (KP4, PaTu8902, MiaPaca and Panc0203) followed by immuno-fluorescence staining confirmed a punctate distribution of MYOF which colocalized with LAMP2 positive lysosomes, with little or no detectable signal on the plasma membrane (Fig. 1h; Supplementary Fig. 1c). In contrast, MYOF-HA localized to the plasma membrane or showed a diffuse cytoplasmic localization in non-PDA cells with no visible overlap with LAMP2-positive structures (Fig. 1h; Supplementary Fig. 1d). To further confirm the differential subcellular targeting of MYOF in PDA versus non-PDA cells we performed biotin labeling of surface proteins in KP4 cells and HEK293T cells transiently expressing MYOF-HA followed by streptavidin-mediated immunoprecipitation. Consistent with the immunofluorescence results, we find that endogenous MYOF is not biotinylated in KP4 cells and therefore absent from the plasma membrane (Fig. 1i). In contrast MYOF transiently expressed in HEK293T cells was efficiently biotinylated, consistent with its plasma membrane localization in these cells (Fig. 1h,i). Thus, Myoferlin is both upregulated and show substantial retargeting to the lysosome in PDA cells. Given the preferential upregulation of MYOF in PDA cells relative to HPDE, we focused our functional analysis on this Ferlin family member in subsequent experiments.

To establish that MYOF resides on the outer (cytoplasm-facing) leaflet of the lysosomal membrane rather than being trafficked to the lysosomal lumen for degradation, we first treated cells with the lysosome V-ATPase inhibitor Bafilomycin A1 (BafA1) to inhibit lysosomal degradation. BafA1 treatment led to a progressive accumulation of known cargo proteins such as the autophagy receptor p62 (Supplementary Fig. 1e). In contrast MYOF did not show an increase in levels upon BafA1 treatment suggesting that it is unlikely to be a cargo protein brought to the lysosome for degradation.

To further prove that MYOF is anchored to the cytoplasmic face of the lysosomal membrane, we treated affinity-captured PDA lysosomes with increasing concentrations of proteinase K, with the expectation that cytoplasmic facing proteins would be sensitive to proteinase K digestion while luminal proteins would be protected. As expected, luminal proteins (LC3, Cathepsin C) and luminal facing proteins (LAMP2) were protected from proteinase K digestion, whereas membrane proteins exposed to the cytoplasm such as NPC1 were progressively digested with increasing concentrations of proteinase K (Fig. 1j). MYOF showed rapid digestion by proteinase K, confirming that it is a membrane protein present on the outer surface of the lysosome (Fig. 1j). Taken together these data indicate that MYOF is topologically anchored via its C-terminal transmembrane domain to the outer lysosomal membrane, with its N-terminus extending into the cytoplasm.

### PDA lysosomes display enhanced protection against membrane damage

Given a known function for Ferlin proteins in maintenance and repair of membrane, we sought to determine whether lysosomal localization of MYOF might confer enhanced protection against membrane damage. To do so, we compared the response of PDA lysosomes versus non-PDA lysosomes under conditions that mimic various forms of lysosomal stress. We first employed acute, chemically-induced lysosomal membrane damage by treating cells with LLOMe (L-Leucyl-L-leucine O-methyl ester), a well-established lysosomotropic agent commonly used to rupture endolysosomal membranes^13, 14, 27^. LLOMe rapidly and efficiently accumulates within the lumen of acidic organelles where it is converted by the lysosomal protease Cathepsin C into a polymer capable of causing small ruptures in the membrane^27^. To test the response of lysosomes to the LLOMe challenge, cells were preloaded with Lysotracker red dye, which accumulates within intact lysosomes, and the rate of dye leakage induced by LLOMe was measured over time. While lysosomes in non-PDA cells (HPDE, U20S and HEK293T) lost virtually all lysotracker red staining within 15min of LLOMe treatment (Fig. 2a; Supplementary Fig. 2a), consistent with progressive lysosomal membrane rupture, PDA lysosomes retained the Lysotracker dye for longer time periods (Fig. 2a,b). This effect was not due to reduced Cathepsin C-dependent activation of LLOMe in PDA lysosomes, as Cathepsin C levels were similar, if not higher in PDA cells relative to HPDE cells (Supplementary Fig. 2b). Prior studies have also established that particulate material such as crystals of silica, alum and uric acid are capable of damaging vesicles and lysosomal membrane^14^. Similarly, changes in osmolarity have also been shown to cause changes in endolysosome membrane tension and permeability^28^. Therefore, we measured the response of PDA and non-PDA cells to these additional forms of endolysosome membrane damage. Consistent with our observations with LLOMe, we find that PDA lysosomes are more resistant to silica-induced and hypertonic sucrose-induced membrane damage as measured by reduced loss of Lysotracker red dye signal (Supplementary Fig. 2c-f). Collectively, these data suggest that, relative to non-PDA counterparts, PDA lysosomes are capable of withstanding diverse membrane perturbing agents, relative to non-PDA lysosomes.

**Fig. 2.**
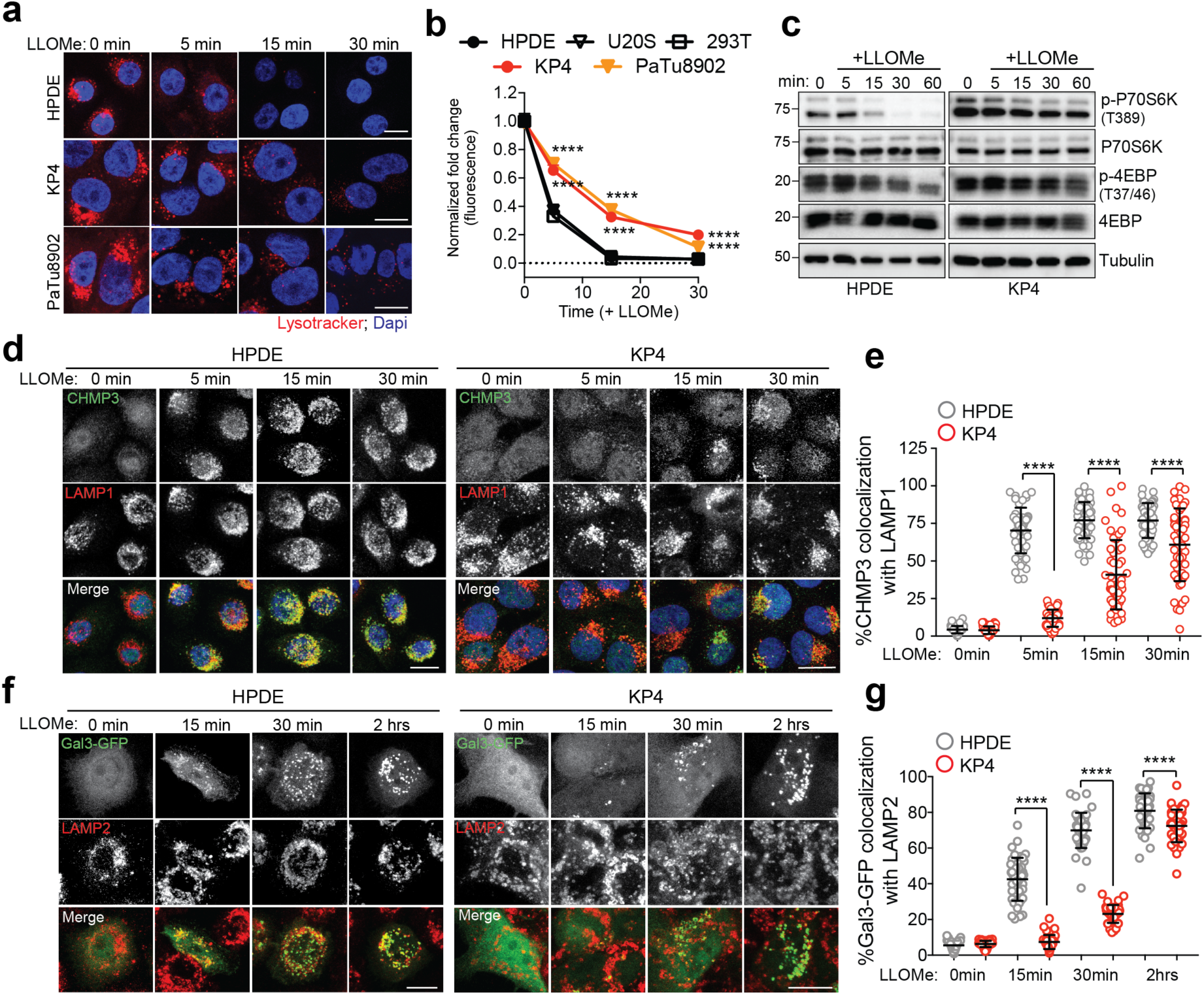
PDA lysosomes are more resistant to lysosome membrane damage. **a.** Time-course of lysotracker red staining in HPDE, KP4 and PaTu8902 following treatment with LLOMe. (HPDE n = 76, 80, 81, 80; KP4 n = 80, 77, 80, 82; PaTu8902 n = 96, 96, 81, 82 cells). **b.** Quantification of normalized fold change in lysotracker red stain in HPDE, U20S, 293T, KP4 and PaTu8902 cells treated with LLOMe. Data are mean ± s.e.m from n = 60-96 cells per timepoint for each cell line. *P* values determined by two-way ANOVA. **c.** Immunoblots for the indicated proteins in HPDE and KP4 cells following time course treatment with LLOMe. **d.** HPDE and KP4 cells with and without LLOMe treatment were co-stained for CHMP3 (green) and LAMP1 (red). **e**. Quantification of percentage co-localization in control and LLOMe treated cells. (HPDE n = 50, 51, 50, 51; KP4 n = 51, 50, 50, 50 cells). **f.** HPDE and KP4 cells expressing GFP-Galectin 3 (green) were treated with LLOMe for the indicated times and co-stained for LAMP2 (red). **g**. Quantification of percentage co-localization in control and LLOMe treated cells. (HPDE n = 39, 43, 40, 42; KP4 n = 43, 42, 40, 42 cells). Scale, 20μm. Data are mean ± s.d. *P* values determined by unpaired two-tailed *t*-tests. **** *P* < 0.0001.

Along with its degradative functions, the lysosome serves as the platform for nutrient signaling via the mechanistic target of rapamycin complex 1 (mTORC1), which must localize to the lysosome surface^1, 19^ and requires an intact lysosomal membrane in order to be activated^16^. Treatment of HPDE cells with LLOMe over a 1h time course led to a rapid decrease in mTORC1 signaling as measured by phosphorylation of downstream targets p70S6K and 4EBP1 (Fig. 2c). In contrast, PDA cells retained mTORC1 signaling activity throughout the 1h LLOMe treatment duration (Fig. 2c; Supplementary Fig. 2g). Similarly, a 4-fold lower dose of LLOMe was sufficient to suppress mTORC1 signaling in HPDE cells, while PDA cells retained mTORC1 signaling even at the highest dose of LLOMe tested (Supplementary Fig. 2h).

Recent studies showed that lysosomal membrane disruption is accompanied by rapid recruitment of ESCRT machinery that facilitates membrane repair^13, 14^. ESCRT proteins are organized into sub-complexes; ESCRT III proteins (CHMPs 1-7 and IST1) are responsible for mediating membrane constriction and fission whereas ALIX (ESCRT II) and TSG101 (ESCRT I) facilitate recruitment of ESCRT III components to the site of damage^13, 14^. Immunofluorescence staining following a time course of LLOMe, Silica or hypertonic sucrose treatment showed that HPDE cells rapidly recruit ALIX to lysosomes post treatment (Supplementary Fig. 3a-c). In contrast, identically treated KP4 cells showed a significant delay in ALIX recruitment (Supplementary Fig. 3a-c). Similarly, LLOMe treatment lead to rapid requirement of ESCRT III proteins CHMP3 and CHMP1A as early as 1min post treatment in HPDE cells and was significantly delayed in PDA cells (Fig. 2d,e; Supplementary Fig. 3d,e). Additional markers of lysosomal membrane damage include the Galectins – cytoplasmic proteins that recognize and bind to inappropriately exposed glycans on the luminal side of lysosomal membrane proteins^15, 17, 29, 30^. Within 15min of LLOMe treatment 45% of LAMP2 positive lysosomes in HPDE cells recruited Galectin 3 (GAL3) (Fig. 2f,g). In contrast, only 8% of lysosomes in PDA cells recruited GAL3, which showed predominantly diffuse staining in PDA cells even after 30min of LLOMe treatment (Fig. 2f,g). To further confirm differential recruitment of GAL3 to lysosomes in response to LLOMe in KP4 and HPDE cells we captured intact lysosomes from LysoTag expressing cells following treatment with LLOMe for 10 minutes. Lysosomes purified from LLOMe treated HPDE cells contained higher levels of GAL3 as measured by immunoblot, compared to lysosomes purified from identically treated KP4 cells (Supplementary Fig. 3f). These findings support the notion that PDA lysosomes display enhanced protection against acute membrane damage compared to their non-PDA counterparts.

### Myoferlin is required to maintain lysosome function in PDA cells

The unique protein composition of PDA lysosomes may help maintain their membrane integrity and function in response to stress. Based on the known membrane repair functions of the Ferlins, we next tested whether lysosomal localization of MYOF could explain the increased stress resistance observed of PDA lysosomes. Suppression of MYOF expression via shRNA mediated knockdown or CRISPR mediated knockout in PDA cell lines led to lysosomes with aberrant morphology and increased diameter (Fig. 3a; Supplementary Fig. 4a-d) – a phenotype commonly associated with lysosome dysfunction^31, 32^. Electron microscopy analysis of PDA cells following knockdown of MYOF confirmed an abundance of enlarged lysosomes which lacked intraluminal content (Fig. 3b). This phenotype is distinct from general inhibition of lysosome digestion following treatment with lysosomotropic agents (Chloroquine), V-ATPase inhibitors (BafA1)^33^ or knockdown of MiT/TFE transcription factors^6^, which commonly result in distended lysosomes filled with undigested, electron-dense content. Consistent with this idea, MYOF deficient lysosomes were unable to accumulate intraluminal Lysotracker dye (Supplementary Fig. 4e). Importantly, MYOF loss triggered a spontaneous lysosomal membrane stress response, associated with constitutive localization of ESCRT proteins ALIX, CHMP3 and CHMP1A to LAMP1 positive lysosomes (Fig. 3c,d; Supplementary Fig. 4f). Together, the alteration in lysosome morphology, lack of Lysotracker dye retention and constitutive recruitment of ESCRT proteins suggests that MYOF serves to protect against lysosome dysfunction in PDA cells. Accordingly, knockdown of MYOF caused a marked accumulation of lipidated LC3B by immunoblotting (Fig. 3e) and increased LC3B-positive puncta by immunofluorescence (Fig. 3f; Supplementary Fig. 4g) as well as delayed clearance of macropinocytosed serum albumin, indicating defective lysosomal proteolysis (Supplementary Fig. 4h). Moreover, loss of MYOF also led to reduced baseline mTORC1 signaling as measured by phosphorylation of p70S6K and 4EBP1, which was suppressed further following treatment with LLOMe (Supplementary Fig. 4i). These data support a novel role for MYOF in maintenance of lysosome function in PDA cells.

**Fig. 3.**
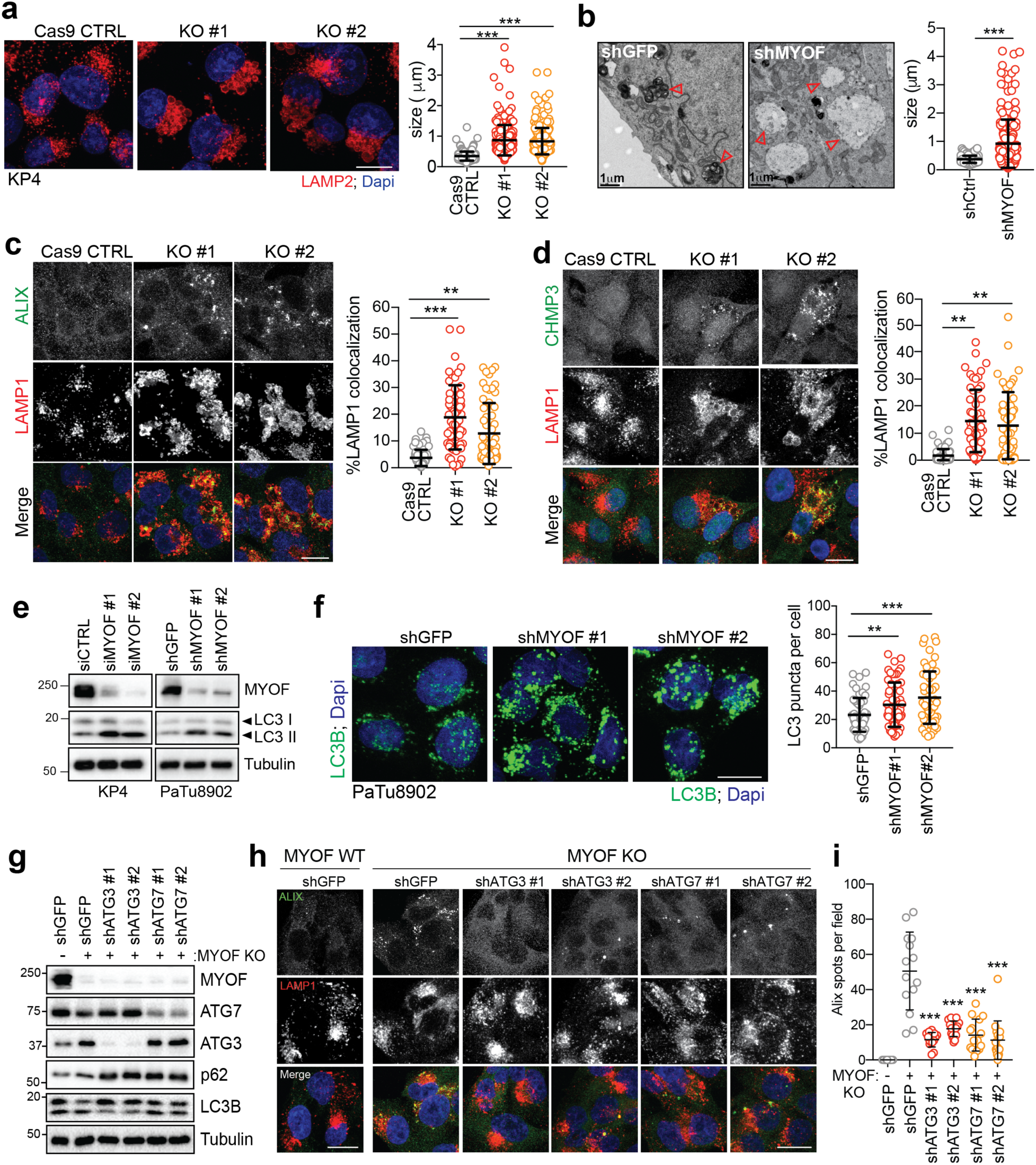
MYOF is essential for lysosome function in PDA cells. **a.** Immunofluorescence staining of LAMP2 (red) following CRISPR mediated knockout of MYOF and Cas9 control KP4 cells. Graph at right shows measurement of lysosome diameter from n = 252 (Cas9 control), 251 (KO#1, KO#2). **b.** Representative electron microscopy images showing aberrant lysosome morphology in KP4 cells following shRNA mediated knockdown of MYOF. Arrowheads highlight differential lysosome morphology in shGFP versus shMYOF conditions. Graph on the right shows quantification of lysosome diameter (n = 200 lysosomes from control; n = 202 lysosomes from MYOF KD cells). Scale bar 1μm. **c, d.** Increased recruitment of ALIX (green) (f) and CHMP3 (green) (g) to LAMP1 positive lysosomes (red) following KO of MYOF in KP4 cells relative to Cas9 control cells. Graphs show the quantification of percentage ALIX (n = 60) and CHMP3 (n = 57) co-localization with LAMP1. **e.** Immunoblot showing increased LC3B lipidation (arrowheads) in KP4 and 8902 cells upon siRNA or shRNA mediated knockdown of MYOF. **f.** Immunofluorescence staining for LC3B (green) showing increase accumulation of LC3B positive autophagosomes in KP4 cells following shRNA mediated knockdown of MYOF compared to control cells. **i.** Graph on the right shows quantification of LC3B puncta (n = 14 fields/condition). **g.** Immunoblot confirming shRNA mediated knockdown of ATG3 and ATG7 and autophagy blockade in KP4 MYOF KO cells. **h.** Recruitment of ALIX (green) to lysosomes (LAMP1; red) in KP4 cells in the presence (WT; n = 14) and absence (KO; n = 14) of MYOF. shRNA mediated knockdown of ATG3 (n = 14, 15) or ATG7 (n = 15, 16) to suppress autophagy causes a decrease in lysosome localization of ALIX in MYOF KO cells. **i.** Graph on the right shows quantification of ALIX puncta per condition. Scale, 20μm. Data are mean ± s.d. *P* values determined by unpaired two-tailed *t*-tests. ** *P* < 0.01; *** *P* < 0.001.

PDA cells are characterized by elevated autophagy and macropinocytosis, which delivers intracellular and extracellular cargo, respectively, to the lysosome for degradation^4, 7, 8^. This increased cargo trafficking may create a unique dependency on MYOF to maintain membrane stability and efficient lysosome function. Therefore, reducing the incoming flux of membrane trafficking should decrease the dependency of PDA cells on MYOF. To explore this idea, we tested whether blocking autophagosome formation and trafficking via knockdown of ATG3 or ATG7 might prevent the lysosomal membrane stress observed in MYOF KO cells. Knockdown of ATG3 or ATG7 in MYOF KO cells led to reduced autophagy as evidenced by accumulation of p62 and unlipidated LC3B-I (Fig. 3g). Strikingly, autophagy suppression resolved the lysosome stress response induced by MYOF loss, as indicated by reduced ALIX recruitment to lysosomes in MYOF KO cells (Fig. 3h,i). These data suggest that the increased vesicle trafficking characteristic of PDA cells imparts heightened stress on the lysosome in order to fuse, process and clear incoming cargo; this stress is counteracted by increased expression and lysosomal targeting of MYOF.

### Inducible lysosome targeting of MYOF is sufficient to protect against acute membrane damage

To gain insights into the mechanism of MYOF-mediated lysosomal stabilization, we tested whether inducible targeting of MYOF to the lysosomal membrane in non-PDA cells would be sufficient to protect against LLOMe-mediated damage. To do so, we used a heterodimerization system (FKBP-FRB) to inducibly recruit a variant of MYOF lacking its C-terminal transmembrane domain (MYOF*Δ*TM) (Fig. 4a), to the lysosome membrane following addition of a rapamycin-derived chemical dimerizer AP21967 (AP)^34, 35^ (Fig. 4b). U2OS cells were first engineered to stably express the FKBP module conjugated to the C-terminus of the lysosomal membrane protein TMEM192 (T192-Flag-FKBP)^34, 35^, which we confirmed localizes to LAMP2 positive lysosomes (Fig. 4c). These cells were then transiently transfected with MYOF*Δ*TM conjugated to FRB* (referred to hereafter as MYOF-FRB*). In the absence of AP, MYOF-FRB* showed a diffuse cytoplasmic localization (Fig. 4d**; top**) in T192-Flag-FKBP expressing U2OS cells. Upon treatment with AP, MYOF-FRB* was massively recruited to LAMP2 positive lysosomes (Fig. 4d**; bottom**). Without AP treatment, addition of LLOMe to U2OS cells led to recruitment of ALIX (Fig. 4e) and GAL3 (Supplementary Fig. 5a) to lysosomes as expected. However, following addition of AP and the resulting recruitment of MYOF-FRB* to lysosomes, recruitment of ALIX and GAL3 to lysosomes was significantly reduced in response to LLOMe (Fig. 4e; Supplementary Fig. 5a). These data suggest that targeting of MYOF to the membrane of non-PDA lysosomes is sufficient to confer protection against chemically induced membrane damage.

**Fig. 4.**
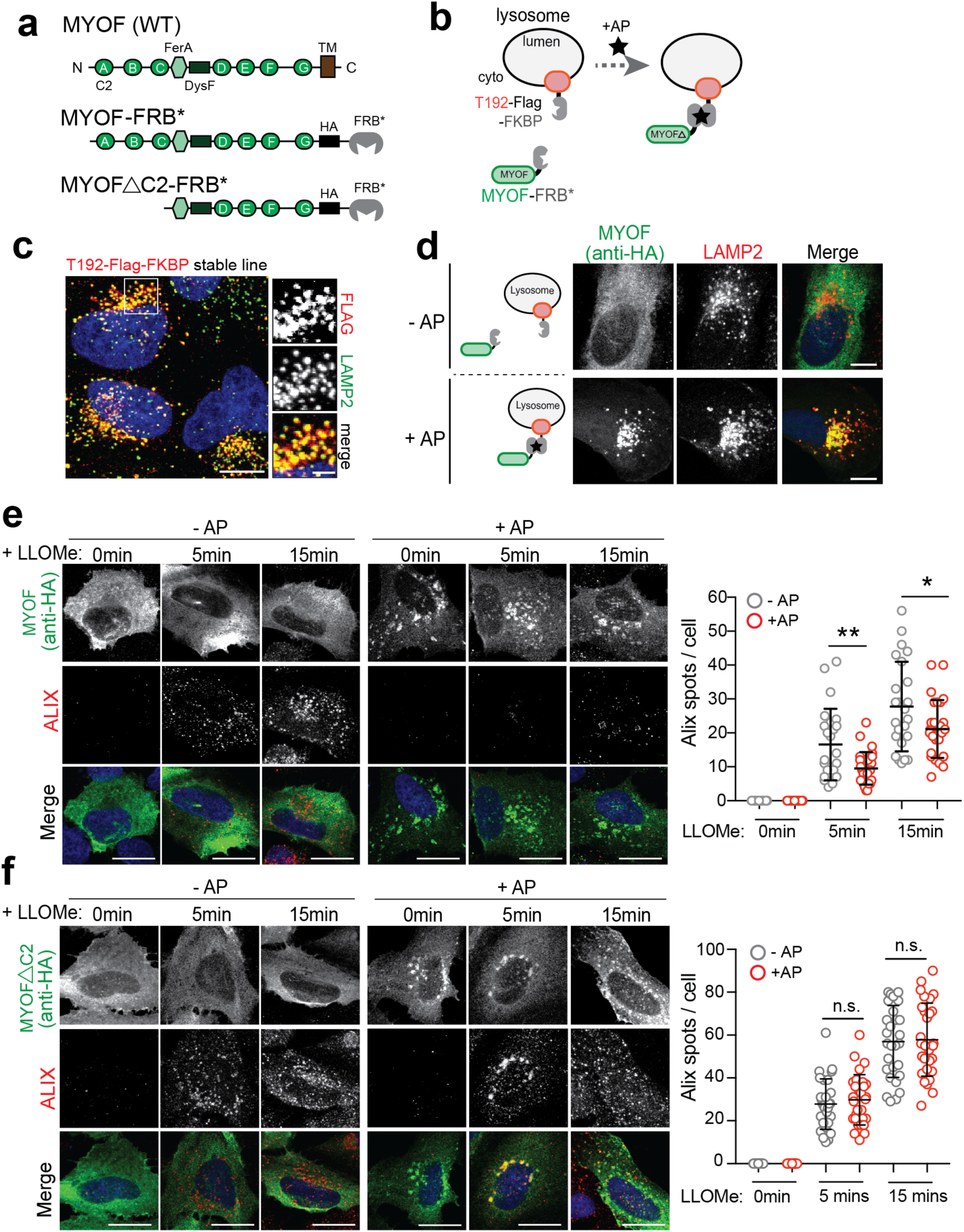
The N terminal C2 domains of MYOF are required for membrane protection. **a.** Domain structure of MYOF and MYOF-FRB* variants. **b.** Schematic showing heterodimerization of MYOF-FRB* to lysosome membrane anchored T192-Flag-FKBP following addition of the rapalogue (AP21967; AP). TM, transmembrane domain. **c.** Lysosomal localization of T192-Flag-FKBP (Flag) in stably expressing U20S cells. **d.** Transient expression of MYOF-FRB* (MYOF; detected with anti-HA antibody) in U20S cells stably expressing T192-Flag-FKBP. In the absence of AP, MYOF-FRB* is cytoplasmic while upon AP addition MYOF*Δ*TM is recruited to LAMP2 positive lysosomes (red). **e.** Recruitment of MYOF-FRB* protects against LLOMe induced damage and ALIX recruitment. U20S-T192-Flag-FKBP expressing cells transfected with MYOF-FRB* were treated with LLOMe for the indicated time points in the absence (n = 30, 24, 24 cells) or presence (n = 30, 27, 27 cells) of AP, followed by immuno-staining for HA (green) and ALIX (red). **f.** U20S-T192-Flag-FKBP cells transfected with MYOF*Δ*C2-FRB were treated as in ‘e’ (-AP n = 29 cells; + AP n = 29 cells) followed by immuno-staining for HA (green) and ALIX (red). Lysosomal recruitment of MYOF*Δ*C2 does not protect against LLOMe induced ALIX recruitment. Graphs at right show quantification of ALIX spots per cell in response to LLOMe. Scale for all panels, 20μm. Data are mean ± s.d. *P* values determined by unpaired two-tailed *t*-tests. * *P* < 0.05; ** *P* < 0.01; n.s. not significant.

The first three C2 domains of MYOF have been shown to bind membrane lipids, alter their distribution^36, 37^ and recruit accessory proteins^38^ and are therefore thought to be key mediators of the membrane resealing functions of MYOF and DYSF. We therefore tested whether a variant of MYOF lacking C2A, C2B and C2C domains (MYOF*Δ*C2) would lead to loss of membrane protection following LLOMe-mediated damage in non-PDA cells. Unlike MYOF-FRB*, AP mediated recruitment of the MYOF*Δ*C2-FRB* mutant to T192-Flag-FKBP positive lysosomes was unable to prevent LLOMe-induced damage, as shown by unchanged ALIX and GAL3 recruitment irrespective of the AP dimerizer (Fig. 4f; Supplementary Fig. 5b). These data establish the N-terminal C2 domains of MYOF as mediating its protective function at the lysosome.

### Loss of MYOF leads to impaired *in vivo* tumour growth

Given the established role for lysosomes as mediators of cellular adaptation and growth in PDA, we next determined the impact of MYOF loss on PDA growth *in vitro* and *in vivo*. Similar to human PDA cell lines (Fig. 1g), tumours isolated from a genetically engineered mouse model of PDA (p48-Cre;LSL-Kras^G12D^;Trp53^L/+^; referred to here as KPC)^39^ displayed levels of Myof that were higher than normal pancreas and liver, and comparable to mouse C2C12 muscle cells (Fig. 5a). Dysf also showed higher levels in KPC tumours relative to normal pancreas (Fig. 5a). Importantly, Myof expression in KPC tumours was restricted to CK19 positive tumour epithelia and did not show significant overlap with *α*-SMA positive stromal cells (Fig. 5b). CRISPR mediated knockout of Myof in murine KPC cells^40^ led to accumulation of LC3B-II, reduced accumulation of Lysotracker dye and reduced *in vivo* growth following transplantation in syngeneic hosts (Supplementary Fig. 6a-d). Similarly, shRNA mediated knockdown or CRISPR mediated knockout of MYOF in KP4 cells significantly reduced colony forming ability (Fig. 5c) and *in vivo* tumour growth (Fig. 5d,e). Resected tumours also showed a decrease in proliferation as measured by Ki67 staining (Fig. 5f). Together these data strongly support a critical role for MYOF as a novel lysosomal membrane protein essential for maintenance of organelle function during tumour growth *in vivo*.

**Fig. 5.**
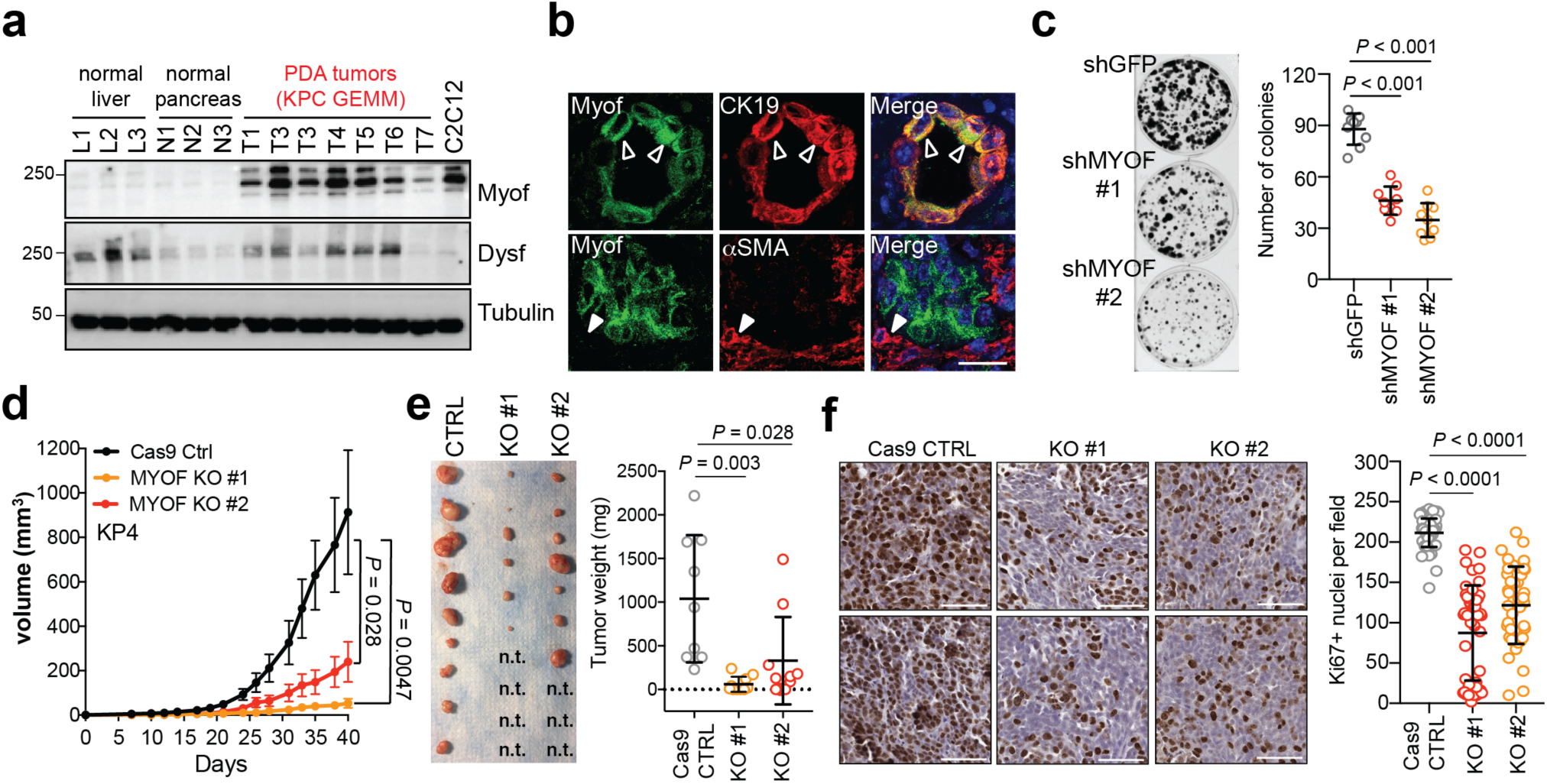
MYOF is required for PDA tumor growth. **a.** Expression levels of Myof and Dysf in mouse KPC derived PDA tumours relative to normal liver, normal pancreas and the mouse myoblast cell line, C2C12. **b.** Immuno-fluorescence staining of Myof (green), the epithelial marker CK19 (red; top) and the stromal marker *α*-SMA (red, bottom) in mouse KPC tumours showing co-localization of Myof with CK19 (open arrowheads) but not *α*-SMA (close arrow heads). Scale, 50μm. **c.** Colony formation of shGFP or shMYOF infected KP4 cells. Graph shows quantification of colony area per condition from n = 9 independent experiments. **d.** *In vivo* growth in nude mice of s.c. KP4 xenografts following CRISPR mediated KO of MYOF. N = 9 (CTRL), 10 (KO#1), 10 (KO#2) tumours per group. Error bars represent s.e.m. **e.** Images (left) and tumour weight (right) of control and MYOF KO KP4 xenografts resected at day 42. n.t. no macroscopic tumour identified upon resection. **f.** Ki67 staining of control and MYOF KO xenografts. Graph shows quantification of Ki67 positive nuclei from n = 56 (control), 46 (KO#1), 46 (KO#2) fields from 3-4 tumours per group. Scale, 100μm.

### MYOF levels dictate worse prognosis in PDA patients

Consistent with high expression levels of MYOF in human PDA cell lines and murine KPC tumours, analysis of several publicly available patient PDA datasets showed that MYOF transcript levels were significantly higher in human PDA tumours relative to normal pancreas or adjacent non-neoplastic tissue (Fig. 6a). In addition, semi-quantitative immunohistochemical analysis revealed significantly increased MYOF protein expression in patient-matched neoplastic versus non-neoplastic epithelium (n = 102, *P <* 2.2×10^-25^) with average histoscores of 3.95 ± 2.3 versus 0.57 ± 0.82, respectively. Further dichotomization (histoscores >5 versus ≤5) revealed high MYOF in 23% of PDA versus 0% of adjacent normal pancreas, (n = 102; *P* < 0.0001) (Fig. 6b,c). No significant associations were noted between high MYOF PDA and clinicopathologic parameters except for a significant correlation with female gender (*P* = 0.008; Supplementary Table 3). Notably, high MYOF PDA was associated with worse overall survival in univariate analysis (hazard ratio = 2.03; 95% CI = 1.29-3.19; Supplementary Table 4a) with a median overall survival of 19.4 months compared to 32 months for patients with low MYOF PDA (log rank *P =* 0.002, Fig. 6d). High MYOF expression was also an independent predicator of worse overall survival in multivariate Cox regression analysis (Supplementary Table 4b). Moreover, a significant correlation between high MYOF expression and worse overall survival was confirmed in an independent PDA patient cohort from The Cancer Genome Atlas (Fig. 6e). Together, analyses from these clinical datasets support a role for MYOF and enhanced lysosome function as predictors of shortened survival in PDA patients.

**Fig. 6.**
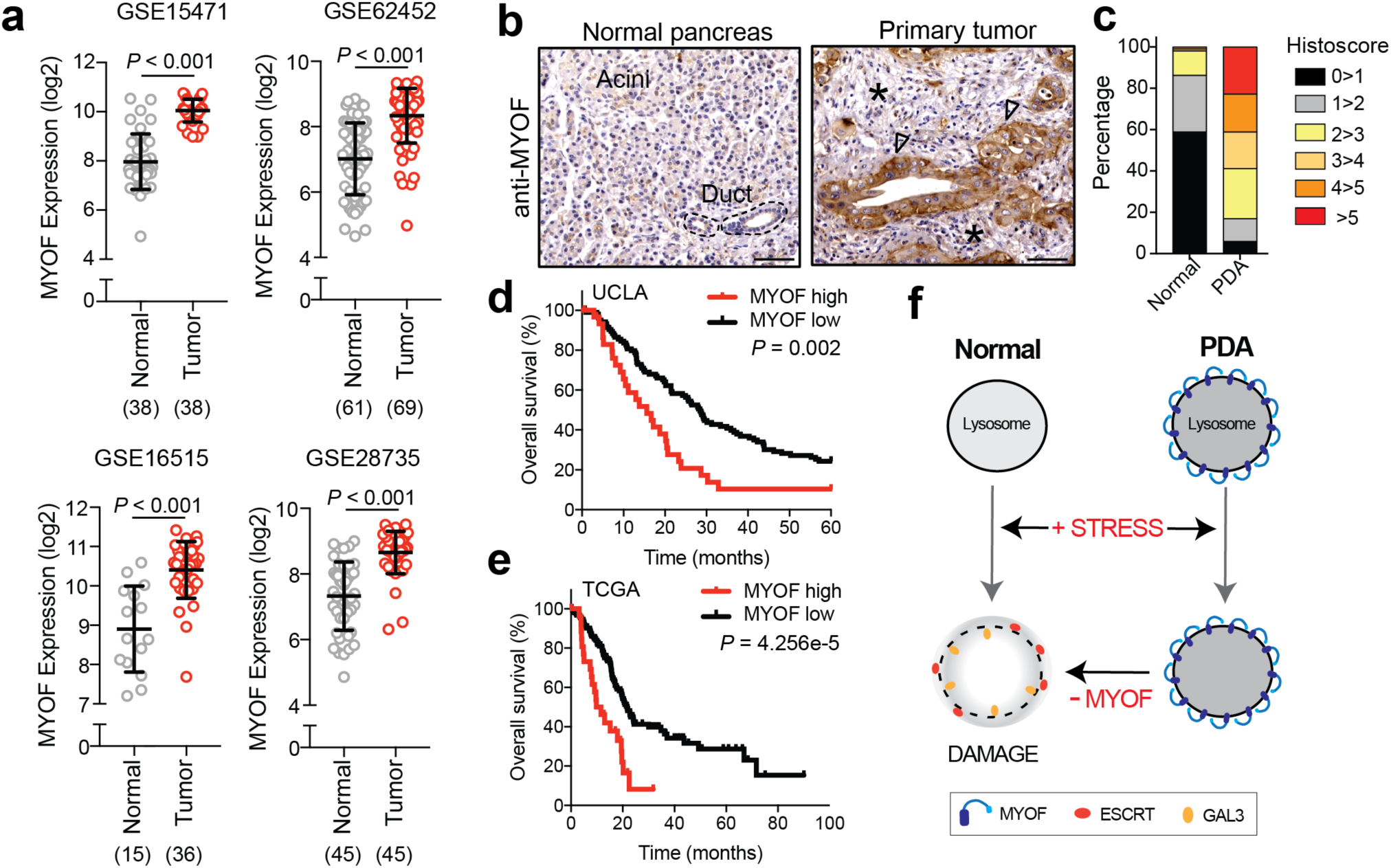
High MYOF expression levels correlate with aggressive disease. **a.** *MYOF* transcript levels in human PDA specimens and normal pancreas (adjacent non-neoplastic tissue) from the indicated datasets. **b.** Immuno-histochemistry showing increased expression of MYOF in primary patient PDA tumour epithelia (arrowheads) compared to normal pancreas or adjacent stroma (asterisk). Scale, 100μm. **c.** Percentage distribution of semi-quantitative histoscore of MYOF staining across normal adjacent (n = 102) and primary PDA (n = 136). **d, e.** High expression of MYOF predicts shorter overall survival in two patient cohorts. N = 136 patients in the UCLA cohort (MYOF high n=31, MYOF low n=105) and n = 185 in The Cancer Genome Atlas (TCGA) cohort (MYOF high; Z score > 1, n = 27; MYOF low Z score < 1, n = 158). p-Value calculated by Log-rank test. **f.** Model comparing lysosomal response to stress in normal (left) and PDA (right) cells. Lysosomal retargeting of MYOF in PDA cells provides protection against membrane stress caused by increased rates of vesicular traffic. Loss of MYOF renders PDA lysosomes more vulnerable to damage. Data are mean ± s.d. *P* values determined by unpaired two-tailed *t*-tests.

## Discussion

Our findings demonstrate that PDA lysosomes are intrinsically more capable of withstanding membrane stress than non-cancer lysosomes. Retargeting of the Ferlin family of plasma membrane repair factors to the lysosomal membrane is critical for this ability and helps to maintain the structural integrity and pro-tumourigenic activities of this organelle. Lysosomal targeting of the Ferlins appears to be an adaptive mechanism in response to the increased degradative burden placed on PDA lysosomes by elevated influx of autophagic and macropinocytic vesicle trafficking (Fig. 6f). Consistent with this model, inhibition of autophagy relieves the lysosomal membrane stress that accumulated following MYOF loss.

The nature of the membrane stress that PDA lysosomes undergo may derive from the specific cargo being degraded, such as large protein aggregates of intracellular or extracellular origin, oxidative damage as well as alterations in membrane composition due to continuous fusion with autophagy-derived and plasma membrane-derived vesicles. Ferlins may counteract this stress through several mechanisms, including formation of oligomers that structurally stabilize the lipid bilayer, thus preventing its rupture, as well as promoting fusion with other vesicles that act as membrane donors to reseal damage that has already occurred. Both mechanisms have been proposed to underlie Ferlin-mediated repair of the plasma membrane of skeletal muscle cells and may operate at the lysosomal membrane in PDA cells as well^21, 23, 24, 37^.

Our findings also suggest that MYOF either replaces or delays activation of a major membrane repair system mediated by the ESCRT complex. ESCRT is thought to repair membrane ‘microtears’ through scission of the free membrane edge into the lumen and resealing of the ‘neck’ driven by subunit polymerization^21, 23, 24, 37^. In turn, this resealing activity is critical to prevent exposure and leakage of luminal lysosomal proteins and protein domains. In the simplest model, the membrane-stabilizing action of MYOF may inhibit or delay ESCRT recruitment by preventing the formation of exposed microtears. Consistent with this idea, loss of MYOF in PDA cells triggers spontaneous ESCRT recruitment to the lysosome. More direct mechanisms of Ferlin mediated ESCRT regulation may also occur, such as competition for a common binding partner. Alternatively, Ferlin proteins may represent a new protective mechanism in cells highly reliant on the lysosome for growth. Given that ESCRT proteins have diverse functions at multiple cellular locations^12^, a dedicated lysosomal program may confer more efficient monitoring and maintenance of lysosomal health.

Accordingly, loss of MYOF leads to severe lysosomal dysfunction and significantly impairs PDA tumour growth *in vivo*, establishing a critical role for this protein in promoting the pro-oncogenic functions of the lysosome in cancer. PDA cells rely on lysosomes as an important source of metabolites^2^ and the lysosome regulates levels of essential micronutrients including iron^41, 42^ and calcium^43^ that can be exchanged with other organelles in the cell including the ER^44, 45^ and mitochondria^41, 42^. Thus, lysosome dysfunction following MYOF loss could indirectly impact additional organelles and cellular metabolic processes. In line with this hypothesis, prior studies^46, 47^ showed that MYOF loss in cancer cells caused respiratory defects in mitochondria, an organelle where MYOF is not found. Finally, high MYOF expression distinguishes a cohort of patient PDA tumours that predict worse overall survival. Several other cancers also show elevated MYOF expression^47–50^ however a role for MYOF in regulation of lysosome function in these cancers remains unknown.

Our findings highlight unique features of cancer lysosomes and identify a dedicated protective mechanism essential for maintenance of lysosomal health. Blocking this protective function of MYOF may pave the way for new lysosome-centered strategies for inhibiting PDA and other cancers.

## Acknowledgements

We thank all the members of the Perera lab and Debnath Lab for helpful discussions. R.M.P is the Nadia’s Gift Foundation Innovator of the Damon Runyon Cancer Research Foundation (DRR-46-17) and is additionally supported by an NIH Director’s New Innovator Award (1DP2CA216364), the Pancreatic Cancer Action Network Career Development Award, and the Hirshberg Foundation for Pancreatic Cancer. H.R.S. is supported by an AACR-Amgen fellowship in Clinical/Translational Cancer Research. R.Z is supported by grants from the NIH (R01GM127763, R01GM130995), a Damon Runyon-Rachleff Innovator Award, Edward Mallinckrodt, Jr. Foundation Grant. D.W.D. receives support from the Hirshberg Foundation for Pancreatic Cancer Research. We thank Reena Zalpuri at the University of California Berkeley Electron Microscope Laboratory for advice and assistance in electron microscopy sample preparation and data collection.

## Author contributions

S.G. performed the majority of experiments and drafted the manuscript. J.Y. developed the FKBP-FRB assay, conducted surface biotinylation experiments, molecular cloning and data analysis. H.H.H. assisted with mouse experiments and immuno-histochemistry. H.R.S. performed the electron microscopy. Z.C. performed the proteinase K protection assay T.I conducted data and pathway analysis. K.W.W and G.K provided pathology analysis of patient samples. R.Z. provided intellectual feedback, reagents and supervised H.R.S. D.W.D provided the PDA TMA and conducted independent pathology analysis and statistical testing. R.M.P. conceived the project, supervised the research, wrote and edited the manuscript.

## Methods

### Cell culture and reagents

The cell lines HPDE, PaTu-8988T, KP4, MiaPaca2, Panc 2.03, PaTu-8902, Panc1, AsPc1, HPAFII, YAPC, Panc 3.27, Panc 10.05, Capan 2 and HupT3 were obtained from the American Type Culture Collection (ATCC) or the DSMZ. Cells were cultured in the following media: PaTu-8988T, KP4, MiaPaca2, PaTu-8902, Panc1, AsPc1, HPAFII, YAPC, Capan2, HEK293T in DMEM supplemented with 10% FBS; Panc 2.03, Panc 3.27, Panc 10.05 and HupT3 in RPMI with 10% FBS; HPDE cells were cultured in keratinocyte serum-free (KSF) medium supplemented by epidermal growth factor and bovine pituitary extract (Life Technologies, Inc., Grand Island, NY). Cell lines were regularly tested and verified to be mycoplasma negative using MycoAlert Detection Kit (Lonza).

FC1245 cells were established from *Pdx1-Cre^+^*, *Kras^LSL-G12D/+^*, *Trp53^lR^*^172^*^H/+^* mice that were backcrossed into a C57BL/6 background and were a gift from David Tuveson (CSHL). Murine PDA cells were maintained in DMEM (Corning) supplemented with 10% FBS (Atlanta Biologicals S11550H) and 1% Pen/Strep (Gibco). Cells were grown in a humidified incubator with 5% CO2 at 37°C. Cultures were routinely verified to be negative for mycoplasma by PCR. Cell lines were authenticated by fingerprinting, and low passage cultures were carefully maintained in a central lab cell bank.

L-leucyl-L-leucine methyl ester (L7393) and Bafilomycin A1 (B1793) were purchased from Sigma and used at 1mM concentration and 100nM respectively. Concentrated stock solutions were prepared in dimethyl sulfoxide (DMSO) and stored at −80 °C in single-use aliquots. Silica nanoparticles (cat.no. tlrl-sio; InvivoGen, San Diego, CA, USA) were suspended in ultrapure water according to manufactures instructions and diluted in complete growth medium to 100 μg/ml. For hypertonic shock, 0.5M sucrose was made in complete growth medium and cells were incubated for the indicated time points. Lysotracker Red-DND-99 (L7528) was purchased from Thermo scientific and used at 75nM. DQ-BSA purchased from Thermo scientific (D12050).

### Constructs

pcDNA3.1-Myoferlin-HA and peGFP-hGalectin-3 was purchased from Addgene (plasmid no. 22443 and 73080 respectively). T192-RFP-3xHA - was generated by subcloning the cDNA of TMEM192 (Origene) together with monomeric Red Fluorescent Protein (mRFP) and 3xHA tag into the Nhe1 and EcoR1 sites of pLJM1 lentiviral vector (addgene plasmid no. 134631) as previously described^9^. To generate MYOFΔTM-HA-FRB*, first an AgeI cut-site was knocked into the pcDNA3.1-Myoferlin-HA plasmid immediately prior to the transmembrane domain by site-directed mutagenesis (Agilent: 210518). Second, a gBlock (IDT) containing HA fused to FRB* was subcloned into the AgeI and KpnI sites of MYOF-HA (Addgene plasmid, #22443) to generate MYOFΔTM-HA-FRB* (referred to as MYOF-FRB*). MYOFΔC2 was generated via PCR amplification from the MYOF-FRB* parent vector. The T192-FLAG-FKBP was a gift from Roberto Zoncu at UC Berkeley.

### shRNAs and siRNAs

shRNA vectors (pLKO.1 puro) were obtained from the Sigma MISSION TRC shRNA library. The sequences and RNAi Consortium clone IDs for the shRNAs used are as follows: shMYOF#1 (human): 5’-GAAAGAGCTGTGCATTATAAA-3’ (TRCN0000320397); shMYOF#2 (human): 5’-GAAAGAGCTGTGCATTATAAA-3’ (TRCN0000001522); shATG3#1 (human): 5’-GATGTGACCATTGACCATATT-3’ (TRCN0000148120); shATG3#2 (human): 5’-GCTGTCATTCCAACAATAGAA-3’ (TRCN0000147381); shATG7#1 (human): 5’-CCCAGCTATTGGAACACTGTA-3’ (TRCN0000007587); shATG7#2 (human): 5’-GCCTGCTGAGGAGCTCTCCAT-3’ (TRCN0000007584); shGFP: 5’-TGCCCGACAACCACTACCTGA-3’ (TRCN0000072186). Pre-designed silencer select siRNAs against MYOF were ordered from Thermo Fischer (Cat# 4392420). shGFP and shScr (Addgene, plasmid #17920) were used as controls.

### Lentiviral-mediated knockdown

For the transfection of lentiviral vectors (pSLIK-hygro, pINDUCER20, and pLKO.1-puro), lentivirus was produced by co-transfection of HEK293T cells with a lentiviral vector and the packaging plasmids psPAX2 (Addgene, plasmid #12260) and pMD2.G (Addgene, plasmid #12259) at a 0.5:0.25:0.25 ratio. Transfection was performed using X-tremeGENE transfection (6365787001; Sigma Aldrich) reagent according to the manufacturer’s instructions. The viral supernatant was collected 48h after transfection and filtered through a 0.45 μm filter. Cells were infected with virus-containing media using Polybrene reagent (TR-1003-G; EMD Millipore) according to the manufacturer’s instructions and selected for 48 hr in 2 μg/mL of puromycin.

### Immunoblotting

Cells were lysed in ice-cold lysis buffer (150 mM NaCl, 20 mM Tris [pH 7.5], 1 mM EDTA, 1 mM EGTA, 1% Triton X-100, 2.5 mM sodium pryophosphate, 1 mM β-glycerophosphate, 1 mM sodium vanadate, and one tablet of Pierce Protease Inhibitor Tablets, EDTA Free [Fisher Scientific-A32965] per 10 mL). Protein content was measured using Pierce BCA Protein Assay Kit (Life Technologies-23227), and 20-30 μg protein was resolved on 8, 10 or 12% protein gels using SDS-PAGE and transferred onto PVDF membranes (EMD MIllipore-IPVH00010). Membranes were blocked in 5% bovine serum albumin (BSA, Sigma Aldrich-A4503) made up in Tris-buffered saline with 0.2% Tween 20 (TBS-T) prior to incubation with primary antibody overnight at 4°C in 5% bovine serum albumin. Membranes were washed in TBS-T and developed after 1h incubation in species-specific horseradish peroxidase-conjugated secondary antibody, visualized using supersignal west pico chemiluminescent substrate (Fisher Scientific-34080), and imaged using the ChemiDoc XRS + System (Biorad).

### Immunoprecipitation

For lysosome immunoprecipitation experiments, human PDA, HPDE or HEK293T cell lines stably expressing T192-mRFP-3xHA were lysed and intact lysosomes from 1–2 mg of total protein was immunoprecipitated using anti-HA-conjugated Dynabeads (Thermo Sci. 88837) as previously described^1, 2^. Proteinase K (Sigma P.2308) digestion was performed when lysosomes were bound to anti-HA-beads. Increasing concentrations of Proteinase K (0.1, 0.5, 1, 2.5ug/ml) in Proteinase K buffer (33.3mM Hepes, 1mM CaCl2, pH:7.4) was added to lysosomes bound to anti-HA beads for 15min on ice. Digestion was terminated with 2mM phenylmethylsulfonyl fluoride (final concentration). Lysosomal proteins were separated by SDS-PAGE and proteins were detected by western blotting.

### Surface Biotinylation assay

The cell surface of KP4 cells and MYOF-HA expressing HEK293T cells was biotinylated on ice using 0.5 mg/ml of EZ-Link™ Sulfo-NHS-SS-Biotin (Thermo Scientific: 21331) for 30min. Cells were washed twice with 20mM Glycine in HBSS on ice to remove excess biotin. Subsequently, cells were lysed in Biotinylation Lysis Buffer (1% Triton X-100, 130mM NaCl, 2.5mM MgCl_2_, 2mM EGTA, 25mM HEPES, pH 7.4, supplemented with protease inhibitor prior to use) on ice for 30min. Lysates were clarified by centrifugation at 13,300 rpm for 10min at 40°C and supernatants were transferred to a new tube. Lysates were quantified by Pierce BCA Protein Assay Kit and equal amounts (∼1mg) of protein were incubated with 100µl of triple-washed Dynabeads™ MyOne™ Streptavidin C1 beads (Invitrogen: 65002) for 2-3h with constant agitation. Beads and captured materials were washed twice in Wash Buffer 1 (2% SDS in dH2O), once in Wash Buffer 2 (0.1% deoxycholate, 1% Triton X-100, 500mM NaCl, 1mM EDTA, and 50mM HEPES, pH 7.5), once in Wash Buffer 3 (250mM LiCl, 0.5% NP-40, 0.5% deoxycholate, 1mM EDTA, and 10mM Tris, pH 8.1), and twice in Wash Buffer 4 (50mM Tris, pH 7.4, and 50mM NaCl). Washes were performed at RT for 5min with gentle agitation. Samples were eluted by boiling in Laemlli buffer.

### FRB-FKBP heterodimerization assay

U2OS cells stably expressing T192-Flag-FKBP were grown on coverslips and transiently transfected with either MYOF-FRB* or MYOFΔC2-FRB* using X-tremegene 9 DNA Transfection Reagent (Roche). After 24h, dimerization was induced by adding 100nM of AP21967 (Takara: 635056) for 1h to cells. Coverslips were then washed twice prior to treated with 1mM LLOME for the indicated timepoints. Cells were subsequently fixed for immunofluorescent staining.

### Immunofluorescence

Human cells were cultured for two days on coverslips coated with fibronectin. After two PBS washes, cells were fixed with paraformaldehyde for 15min at room temperature or with ice-cold methanol for 5min at -20^0^C. PFA fixed cells were permeabilized with 0.1% Saponin or 0.3% Triton X-100 for 5min. Samples were then blocked with 5% normal goat serum for 15min at room temperature prior to incubation with primary antibodies overnight at 4^0^C (Supplementary Table 1). After washing three times with PBS, cells were incubated in secondary antibody (diluted 1:400 in PBS) at room temperature for 30min. Slides were mounted on glass slides using DAPI Fluoromount-G (0100-20, SouthernBiotech) and imaged on a Zeiss Laser Scanning Microscope (LSM) 710 using a 63x objective. Image processing and quantification were performed using ImageJ. For DQ-BSA experiments, cells were incubated with 200μg DQ-BSA for 3h and subsequently chased for 3h in DQ-BSA free media. Where indicated, treatment with BafA1 was for 1h at 100nM. The number of DQ-BSA spots co-localizing with LAMP2 positive lysosomes was quantified. For experiments requiring exposure to LLOMe, silica nanoparticles, hypertonic sucrose or Lysotracker, coverslips were bathed in medium containing the appropriate reagent as indicated, prior to fixing and staining. Measurement of colocalization was conducted using thresholded images using the image calculator function in Image J. Overlapping pixels in the red and green channels were measured and percentage overlap calculated from a minimal of 10 fields per condition. For measuring total cell fluorescence, analysis was performed on a per cell basis using Image J. Mean fluorescence intensities for each cell was determined by subtracting the mean fluorescence of the background in each image.

### Generation of Myoferlin knockout cell lines using CRISPR/Cas9

Myoferlin knockouts in KP4, Patu8902 and FC1245 cells were generated using the RNP-electroporation method as previously described (Liang et al., 2015). One million cells were electroporated using the Amaxa 4D Nucleofector kit (V4XC-9064, Lonza). Guide RNA and Cas9 complexes were pre-formed using 160μM crRNA annealed to 160μM tracrRNA (Dharmacon) and incubated with 40μM Cas9 protein (purchased from University of California, Berkeley). Cutting efficiency was assessed 48h post-electroporation using PCR and sanger sequencing. Myoferlin knockout was confirmed using quantitative RT-PCR and immunoblotting after clonal expansion of single cells.

Guide RNA Sequences (5’-3’):

Mouse Myoferlin exon 3 - TGACTT GAG GGG GAT ACC AC

Mouse Myoferlin exon 4 - CTC CCT GAA GGA CCT GAT TG

Human MYOFERLIN exon 1 – TTT CGT TTT AGG GAT ATT GC

Human MYOFERLIN exon 3 – ATT TTG GAG TTT GAC TTG AG

PCR Primer Sequences (5’-3’):

Ms Myof Ex3 Fwd – tgagtcagaggtttggtgaccc

Ms Myof Ex3 Rev – cagactcgtgacggctgagtat

Ms Myof Ex4 Fwd – agccaaagagaggagcatgtgt

Ms Myof Ex4 Rev – tatctcaacctcccaactgccg

Hu MYOF Ex1 Fwd – GGGAGTTCGGTATCAGTTTACA

Hu MYOF Ex1 Rev – CTGGAGAGACTTGGCTTCATC

Hu MYOF Ex 3 Fwd – TCAGCTGCCTTCAGGTTTAG

Hu MYOF Ex3 Rev – CCACATCTGCTATTGGCTTAGA

### Histology and immunostaining

Tissue samples were fixed overnight in 10% formalin, and then embedded in paraffin and sectioned (5mm thickness) by the UCSF mouse histopathology core. Haematoxylin and eosin staining was performed using standard methods. Slides were baked at 60°C for a 1h, deparaffinized in xylenes (three treatments, 5 min each), rehydrated sequentially in ethanol (5min in 100%, 5min in 90%, 5min in 70%, 5min in 50%, and 5min in 30%), and washed for 5min in water twice. For antigen unmasking, specimens were cooked in a 10mM sodium citrate buffer (pH 6.0) for 10min at 95°C using conventional pressure cooker, rinsed three times with PBS, incubated for 1h with 3% H_2_O_2_ at room temperature to block endogenous peroxidase activity, washed three times with PBS, and blocked with 2.5% goat serum in PBS for 1h. Primary antibodies were diluted in blocking solution and incubated with the tissue sections at 4°C overnight. Specimens were then washed three times for 5min each in PBS and incubated with secondary anti-mouse/rabbit IgG (Vector Laboratories, MP-7500) or fluorescent-conjugated secondary antibodies (1:500, Supplementary Table 1) at RT for 1h. Following three washes in PBS, slides were stained for peroxidase for 3min with the DAB (di-aminebenzidine) substrate kit (SK-4100, Vector Laboratories), washed with water and counterstained with haematoxylin or mounted on glass slides using DAPI Fluoromount-G. Bright light images were obtained with a with a KEYENCE BZ-X710 microscope. Fluorescent images were obtained with Zeiss Laser Scanning Microsope (LSM) 710.

### Human Samples

The pancreatic cancer tissue microarray has been previously described^3^. Clinicopathologic variables used for analysis of the TMA were based on the 7^th^ edition of the AJCC/UICC TNM staging system for pancreatic cancer. Immunohistochemical staining for MYOF was conducted using anti-MYOF ab (Sigma; Cat. No HPA014245). Multiple (2-3) 1.0 mm cores for each tumour in the TMA were independently scored for staining positivity by two pathologists (DWD and KWW) using modified histoscores (range 0-9), representing the product of staining intensity (0-3, 0- absent; 1-weak; 2-moderate; 3-strong) and percentage of tumour cell staining (0-3, 0, none; 1, 1-33%; 2, 34-66%; 3, 67-100%). For any core where histoscore differed by more than 2 between the two observers, a revised score was assigned based on consensus evaluation. The average histoscore for both observers was used for subsequent analysis. For dichotomization, each tumour was assigned to either a low (histoscore 0-5) or high (histoscore >5-9) staining group. Survival estimates were generated using the Kaplan-Meier method and compared using log-rank tests. Multivariate Cox proportional hazards models were used to test statistical independence and significance of multiple predictors with backward selection performed using the Akaike Information Criterion. Overall survival time was measured from the date of surgery to the date of death due to any cause or last clinical visit up to 60 months.

PDAC expression data was collected from the following databases: GSE62452, GSE28735, GSE15471, GSE16515, GSE43795. *P* values were calculated by paired or unpaired, two-tailed *t-* test.

### Animal Experiments

All experiments were carried out in a clean conventional facility. For subcutaneous xenografts, 2 million KP4 Cas9 control/MYOF KO or FC1245 Cas9 control/myof KO cells were injected into the flank of nude mice and C57Bl/6 mice respectively (5-6 mice per group). Tumour length and width were measured thrice a week and the volume was calculated according to the formula (length x width^2^)/2. Mice were euthanized when tumour volume reached 800-1000mm^3^ and tumours were harvested and submitted for histological examination. Mouse strains were obtained from Jackson Laboratories and kindly provided by colleagues: Ptf1a-Cre (Jax stock no. 019378) mice from C. Wright, LSL-Kras^G12D^ (Jax stock no. 008179) mice from D. Tuveson and T. Jacks and Tp53^Lox/Lox^ (Jax stock no. 008462) mice from A. Berns.

### Transmission electron microscopy

KP4 cells (control and Myoferlin knockdown cells) grown on tissue culture plates were fixed in 2.5% glutaraldehyde 0.1M sodium cacodylate buffer, pH 7.4 (EMS, Hatfield, PA, USA) for 30min at room temperature and then gently scraped using a rubber policeman. Cell pellets were stabilized in 1% low melting point agarose, then cut into 1mm cubes. Samples were rinsed (3x; 10min, RT) in 0.1M sodium cacodylate buffer, pH 7.2, and immersed in 1% osmium tetroxide with 1.6% potassium ferricyanide in 0.1M sodium cacodylate buffer for 1 hour. Samples were rinsed (3x; 10min, RT) in buffer and then in distilled water (3x; 10min, RT). Sample were immersed in 0.5% aqueous uranyl acetate *en bloc* stain for 1h, protected from light and then rinsed in water (3x; 10min, RT). Samples were then subjected to an ascending acetone gradient (10min; 35%, 50%, 70%, 80%, 90%) followed by pure acetone (2x; 10min, RT). Samples were progressively infiltrated while rocking with Epon resin (EMS, Hatfield, PA, USA) and polymerized at 60 ^0^C for 24-48h. Thin sections (70nm) were cut using a Reichert Ultracut E (Leica, Wetzlar, Germany). Sections were then collected onto formvar-coated 200 mesh copper grids. The grids were post-stained with 2% uranyl acetate followed by Reynold’s lead citrate, for 5min each. The sections were imaged using a Tecnai 12 120kV TEM (FEI, Hillsboro, OR, USA) and data recorded using an UltraScan 1000 with Digital Micrograph 3 software (Gatan Inc., Pleasanton, CA, USA).

### Pathway analysis and statistical analysis

A list of *³*2 fold significantly enriched (p-value *£*0.05; Log_2_ fold change *³* 1) proteins present in PDA lysosome elutes was analyzed for significantly overrepresented gene ontology (GO) terms (Biological Processes) and biological pathways (KEGG) using g:Profiler g:GO St tool^4^ and The Database for Annotation, Visualization and Integrated Discovery (DAVID).

Experimental data were analyzed using GraphPad Prism and excel built-in tests and are indicated in the figure legends. For all graphs, error bars indicate mean ± standard deviation unless otherwise indicated. Numbers of samples analyzed per experiment are reported in the respective figure legends.

## Supplementary Tables

**Supplementary Table 1:** ≥2 fold significantly enriched proteins identified in PDA lysosome elutes.

| Accession Number | avg<br>HEK293T | avg<br>PaTu8988T | fold<br>(PaTu8988T/<br>HEK293T) | ttest | log2<br>fold<br>change | (-log10<br>ttest) |
| --- | --- | --- | --- | --- | --- | --- |
| NP_001123927 | 1 | 89.66667 | 89.66667 | 7.1E-05 | 6.486 | 4.146 |
| NP_002517 | 1 | 68.66667 | 68.66667 | 6.5E-06 | 6.102 | 5.188 |
| NP_002195 | 1 | 52.33333 | 52.33333 | 1E-05 | 5.71 | 4.993 |
| NP_079105 | 1 | 50.66667 | 50.66667 | 1.2E-05 | 5.663 | 4.936 |
| XP_005264575 | 0.66667 | 23.66667 | 35.5 | 0.00014 | 5.15 | 3.868 |
| XP_005264575 | 0.66667 | 23.66667 | 35.5 | 0.00014 | 5.15 | 3.868 |
| NP_002107 | 1 | 34.66667 | 34.66667 | 0.00025 | 5.115 | 3.606 |
| NP_001290453 | 1 | 33.66667 | 33.66667 | 8.7E-05 | 5.073 | 4.061 |
| NP_001093640 | 1 | 32 | 32 | 1.1E-05 | 5 | 4.941 |
| NP_001193773 | 1 | 31.66667 | 31.66667 | 0.00096 | 4.985 | 3.016 |
| XP_016856023 | 1 | 31.33333 | 31.33333 | 3.1E-05 | 4.97 | 4.507 |
| XP_011543028 | 1 | 31 | 31 | 0.00013 | 4.954 | 3.871 |
| NP_038479 | 3.66667 | 108 | 29.45455 | 3E-07 | 4.88 | 6.52 |
| NP_038479 | 3.66667 | 108 | 29.45455 | 3E-07 | 4.88 | 6.52 |
| NP_001073286 | 1 | 27.66667 | 27.66667 | 5.2E-05 | 4.79 | 4.285 |
| NP_000201 | 1 | 27 | 27 | 2.3E-05 | 4.755 | 4.638 |
| XP_006711978 | 1 | 26.66667 | 26.66667 | 0.00039 | 4.737 | 3.411 |
| XP_016874073 | 1 | 26.33333 | 26.33333 | 4.5E-05 | 4.719 | 4.345 |
| XP_016874073 | 1 | 26.33333 | 26.33333 | 4.5E-05 | 4.719 | 4.345 |
| NP_001073286 | 1 | 26.33333 | 26.33333 | 0.00014 | 4.719 | 3.865 |
| XP_005266132 | 1 | 26 | 26 | 0.00041 | 4.7 | 3.384 |
| NP_004159 | 1 | 25 | 25 | 0.0024 | 4.644 | 2.62 |
| NP_004159 | 1 | 25 | 25 | 0.0024 | 4.644 | 2.62 |
| XP_011514006 | 1 | 24 | 24 | 0.00134 | 4.585 | 2.871 |
| NP_001007069 | 1 | 23.33333 | 23.33333 | 4.7E-06 | 4.544 | 5.325 |
| NP_000253 | 1 | 22.66667 | 22.66667 | 1.6E-05 | 4.503 | 4.788 |
| NP_009220 | 1 | 22.33333 | 22.33333 | 3.6E-07 | 4.481 | 6.447 |
| NP_001007069 | 1 | 22.33333 | 22.33333 | 5.7E-06 | 4.481 | 5.245 |
| NP_000593 | 1 | 21.66667 | 21.66667 | 2E-05 | 4.437 | 4.706 |
| NP_001229971 | 1 | 21.33333 | 21.33333 | 0.00236 | 4.415 | 2.627 |
| NP_001007070 | 1 | 21 | 21 | 4.1E-06 | 4.392 | 5.383 |
| NP_001035861 | 1 | 20.66667 | 20.66667 | 0.00037 | 4.369 | 3.434 |
| XP_016883302 | 1 | 18.33333 | 18.33333 | 0.00013 | 4.196 | 3.872 |
| XP_011531200 | 1 | 17.66667 | 17.66667 | 4.6E-05 | 4.143 | 4.336 |
| NP_001157789 | 0.66667 | 11 | 16.5 | 0.00344 | 4.044 | 2.463 |
| NP_001257904 | 0.66667 | 10.66667 | 16 | 0.0019 | 4 | 2.722 |
| NP_005403 | 0.66667 | 10.33333 | 15.5 | 0.00292 | 3.954 | 2.535 |
| NP_005403 | 0.66667 | 10.33333 | 15.5 | 0.00292 | 3.954 | 2.535 |
| NP_037364 | 1 | 15.33333 | 15.33333 | 0.00356 | 3.939 | 2.448 |
| NP_005219 | 4.66667 | 70.33333 | 15.07143 | 0.00012 | 3.914 | 3.922 |
| NP_005219 | 4.66667 | 70.33333 | 15.07143 | 0.00012 | 3.914 | 3.922 |
| NP_443164 | 1 | 14.66667 | 14.66667 | 0.00149 | 3.874 | 2.825 |
| NP_057231 | 0.66667 | 9.66667 | 14.5 | 0.00067 | 3.858 | 3.172 |
| NP_001679 | 0.66667 | 9.66667 | 14.5 | 0.00882 | 3.858 | 2.054 |
| NP_065843 | 1 | 14.33333 | 14.33333 | 0.00038 | 3.841 | 3.425 |
| NP_001247419 | 1 | 13.66667 | 13.66667 | 0.00199 | 3.773 | 2.701 |
| NP_000916 | 1 | 13.66667 | 13.66667 | 0.00199 | 3.773 | 2.701 |
| XP_016856023 | 2.66667 | 35 | 13.125 | 2.4E-05 | 3.714 | 4.617 |
| NP_001611 | 8.66667 | 112 | 12.92308 | 0.00141 | 3.692 | 2.851 |
| NP_001611 | 8.66667 | 112 | 12.92308 | 0.00141 | 3.692 | 2.851 |
| NP_003557 | 2.33333 | 29.66667 | 12.71429 | 0.00057 | 3.668 | 3.241 |
| NP_060530 | 1 | 12.66667 | 12.66667 | 4E-06 | 3.663 | 5.4 |
| NP_001020527 | 1 | 12.66667 | 12.66667 | 0.00063 | 3.663 | 3.2 |
| NP_001020527 | 1 | 12.66667 | 12.66667 | 0.00063 | 3.663 | 3.2 |
| NP_004915 | 5.66667 | 71 | 12.52941 | 3.2E-06 | 3.647 | 5.499 |
| XP_011528984 | 1.33333 | 16.33333 | 12.25 | 0.00279 | 3.615 | 2.555 |
| NP_003253 | 0.66667 | 8 | 12 | 0.00114 | 3.585 | 2.942 |
| NP_006283 | 1 | 12 | 12 | 0.00197 | 3.585 | 2.705 |
| NP_001171114 | 0.66667 | 8 | 12 | 0.01169 | 3.585 | 1.932 |
| NP_000518 | 1.33333 | 15.66667 | 11.75 | 0.00065 | 3.555 | 3.184 |
| NP_004915 | 4.66667 | 53.33333 | 11.42857 | 1.9E-05 | 3.515 | 4.725 |
| NP_001690 | 1 | 11.33333 | 11.33333 | 0.00101 | 3.503 | 2.998 |
| NP_000174 | 1 | 11.33333 | 11.33333 | 0.00272 | 3.503 | 2.566 |
| NP_000174 | 1 | 11.33333 | 11.33333 | 0.00272 | 3.503 | 2.566 |
| XP_006718556 | 1 | 11 | 11 | 0.00056 | 3.459 | 3.25 |
| NP_079472 | 0.66667 | 7.333333 | 11 | 0.00086 | 3.459 | 3.063 |
| NP_001017423 | 1.33333 | 14.33333 | 10.75 | 0.00124 | 3.426 | 2.905 |
| NP_001093 | 3.66667 | 39 | 10.63636 | 2.9E-05 | 3.411 | 4.534 |
| NP_891988 | 4.33333 | 45.66667 | 10.53846 | 3.1E-06 | 3.398 | 5.515 |
| XP_006717309 | 2.66667 | 28 | 10.5 | 1.8E-05 | 3.392 | 4.75 |
| NP_003840 | 1.33333 | 14 | 10.5 | 0.00199 | 3.392 | 2.701 |
| NP_570846 | 1 | 10.33333 | 10.33333 | 0.01001 | 3.369 | 2 |
| NP_006491 | 1 | 10 | 10 | 9.9E-05 | 3.322 | 4.005 |
| NP_006491 | 1 | 10 | 10 | 9.9E-05 | 3.322 | 4.005 |
| NP_079199 | 1.66667 | 16.33333 | 9.8 | 0.00204 | 3.293 | 2.691 |
| NP_079199 | 1.66667 | 16.33333 | 9.8 | 0.00204 | 3.293 | 2.691 |
| NP_000010 | 1.66667 | 16 | 9.6 | 0.00579 | 3.263 | 2.237 |
| NP_006102 | 1 | 9.333333 | 9.333333 | 0.0007 | 3.222 | 3.155 |
| NP_000681 | 1 | 9.333333 | 9.333333 | 0.00227 | 3.222 | 2.644 |
| NP_001136336 | 0.66667 | 6 | 9 | 8.9E-05 | 3.17 | 4.05 |
| XP_011512751 | 1 | 9 | 9 | 0.00228 | 3.17 | 2.642 |
| NP_003970 | 1 | 9 | 9 | 0.00228 | 3.17 | 2.642 |
| NP_001289561 | 1 | 9 | 9 | 0.00635 | 3.17 | 2.197 |
| NP_001289561 | 1 | 9 | 9 | 0.00635 | 3.17 | 2.197 |
| NP_001034579 | 1 | 9 | 9 | 0.01613 | 3.17 | 1.792 |
| NP_004484 | 0.66667 | 6 | 9 | 0.0329 | 3.17 | 1.483 |
| NP_001449 | 4 | 35.33333 | 8.833333 | 0.00036 | 3.143 | 3.447 |
| XP_005248242 | 1 | 8.666667 | 8.666667 | 0.00033 | 3.115 | 3.486 |
| NP_000134 | 1 | 8.666667 | 8.666667 | 0.0031 | 3.115 | 2.509 |
| NP_071436 | 1 | 8.333333 | 8.333333 | 0.00039 | 3.059 | 3.411 |
| NP_002626 | 1 | 8.333333 | 8.333333 | 0.01417 | 3.059 | 1.849 |
| NP_000204 | 1 | 8.333333 | 8.333333 | 0.0168 | 3.059 | 1.775 |
| NP_000204 | 1 | 8.333333 | 8.333333 | 0.0168 | 3.059 | 1.775 |
| NP_036475 | 1 | 8.333333 | 8.333333 | 0.02844 | 3.059 | 1.546 |
| NP_000970 | 1.33333 | 11 | 8.25 | 0.00192 | 3.044 | 2.716 |
| XP_016869431 | 4.33333 | 35.66667 | 8.230769 | 7.7E-05 | 3.041 | 4.114 |
| NP_037506 | 4.66667 | 38.33333 | 8.214286 | 0.00518 | 3.038 | 2.286 |
| NP_037506 | 4.66667 | 38.33333 | 8.214286 | 0.00518 | 3.038 | 2.286 |
| NP_002831 | 2.66667 | 21.66667 | 8.125 | 2.3E-06 | 3.022 | 5.645 |
| NP_009055 | 1 | 8 | 8 | 0.00027 | 3 | 3.576 |
| NP_009055 | 1 | 8 | 8 | 0.00027 | 3 | 3.576 |
| NP_955394 | 2 | 16 | 8 | 0.00374 | 3 | 2.427 |
| NP_001309906 | 1 | 8 | 8 | 0.00374 | 3 | 2.427 |
| NP_001309906 | 1 | 8 | 8 | 0.00374 | 3 | 2.427 |
| NP_001186913 | 1 | 8 | 8 | 0.01016 | 3 | 1.993 |
| NP_001688 | 1 | 8 | 8 | 0.0249 | 3 | 1.604 |
| NP_955394 | 2 | 15.66667 | 7.833333 | 0.00378 | 2.97 | 2.423 |
| NP_001001937 | 7.33333 | 57 | 7.772727 | 0.00071 | 2.958 | 3.151 |
| NP_001310245 | 1 | 7.666667 | 7.666667 | 0.00056 | 2.939 | 3.25 |
| NP_001310245 | 1 | 7.666667 | 7.666667 | 0.00056 | 2.939 | 3.25 |
| NP_061329 | 1 | 7.666667 | 7.666667 | 0.00749 | 2.939 | 2.125 |
| NP_001177910 | 0.66667 | 5 | 7.5 | 0.00797 | 2.907 | 2.099 |
| NP_001177910 | 0.66667 | 5 | 7.5 | 0.00797 | 2.907 | 2.099 |
| NP_056991 | 1 | 7.333333 | 7.333333 | 0.00199 | 2.874 | 2.701 |
| XP_016881019 | 1 | 7.333333 | 7.333333 | 0.00897 | 2.874 | 2.047 |
| XP_016881019 | 1 | 7.333333 | 7.333333 | 0.00897 | 2.874 | 2.047 |
| XP_006716370 | 1 | 7.333333 | 7.333333 | 0.0191 | 2.874 | 1.719 |
| NP_000173 | 2.33333 | 17 | 7.285714 | 0.00446 | 2.865 | 2.351 |
| NP_001230209 | 2.33333 | 17 | 7.285714 | 0.00579 | 2.865 | 2.238 |
| NP_001089 | 1 | 7 | 7 | 0.00048 | 2.807 | 3.315 |
| NP_000681 | 1 | 7 | 7 | 0.00388 | 2.807 | 2.411 |
| NP_003356 | 1 | 7 | 7 | 0.01324 | 2.807 | 1.878 |
| NP_004386 | 1 | 7 | 7 | 0.03032 | 2.807 | 1.518 |
| NP_490647 | 3.66667 | 24.66667 | 6.727273 | 0.01092 | 2.75 | 1.962 |
| NP_055635 | 1 | 6.66667 | 6.66667 | 7E-05 | 2.737 | 4.154 |
| NP_003357 | 1 | 6.66667 | 6.66667 | 0.00105 | 2.737 | 2.979 |
| NP_005551 | 1 | 6.66667 | 6.66667 | 0.00302 | 2.737 | 2.521 |
| NP_005994 | 1 | 6.66667 | 6.66667 | 0.00921 | 2.737 | 2.036 |
| NP_005995 | 1 | 6.66667 | 6.66667 | 0.03251 | 2.737 | 1.488 |
| NP_005995 | 1 | 6.66667 | 6.66667 | 0.03251 | 2.737 | 1.488 |
| NP_066925 | 2.66667 | 17.66667 | 6.625 | 0.0004 | 2.728 | 3.398 |
| NP_490647 | 3.66667 | 24 | 6.545455 | 0.01403 | 2.71 | 1.853 |
| NP_002831 | 3.33333 | 21.66667 | 6.5 | 4.1E-05 | 2.7 | 4.385 |
| NP_002380 | 0.66667 | 4.333333 | 6.5 | 0.00793 | 2.7 | 2.101 |
| NP_076997 | 0.66667 | 4.333333 | 6.5 | 0.02947 | 2.7 | 1.531 |
| NP_004433 | 1 | 6.333333 | 6.333333 | 8.9E-05 | 2.663 | 4.05 |
| XP_016873572 | 1 | 6.333333 | 6.333333 | 0.00132 | 2.663 | 2.878 |
| NP_002834 | 1 | 6.333333 | 6.333333 | 0.00132 | 2.663 | 2.878 |
| NP_005594 | 1 | 6.333333 | 6.333333 | 0.00377 | 2.663 | 2.423 |
| NP_001922 | 1 | 6.333333 | 6.333333 | 0.01135 | 2.663 | 1.945 |
| NP_002283 | 1 | 6.333333 | 6.333333 | 0.01613 | 2.663 | 1.792 |
| NP_038470 | 1.33333 | 8.333333 | 6.25 | 0.00702 | 2.644 | 2.154 |
| NP_038470 | 1.33333 | 8.333333 | 6.25 | 0.00702 | 2.644 | 2.154 |
| NP_057494 | 0.66667 | 4 | 6 | 0.00056 | 2.585 | 3.25 |
| XP_006713173 | 0.66667 | 4 | 6 | 0.00749 | 2.585 | 2.125 |
| XP_005273672 | 1 | 6 | 6 | 0.01235 | 2.585 | 1.908 |
| NP_001139433 | 4.66667 | 27 | 5.785714 | 2.9E-05 | 2.532 | 4.533 |
| NP_001291395 | 1 | 5.66667 | 5.66667 | 0.00015 | 2.503 | 3.821 |
| NP_003349 | 1 | 5.66667 | 5.66667 | 0.00015 | 2.503 | 3.821 |
| NP_001157666 | 1 | 5.66667 | 5.66667 | 0.00219 | 2.503 | 2.659 |
| NP_006280 | 1 | 5.66667 | 5.66667 | 0.00612 | 2.503 | 2.213 |
| NP_006280 | 1 | 5.66667 | 5.66667 | 0.00612 | 2.503 | 2.213 |
| NP_001317939 | 1 | 5.66667 | 5.66667 | 0.04055 | 2.503 | 1.392 |
| NP_001317939 | 1 | 5.66667 | 5.66667 | 0.04055 | 2.503 | 1.392 |
| NP_001677 | 11 | 62 | 5.636364 | 2.2E-05 | 2.495 | 4.66 |
| NP_004068 | 2 | 11 | 5.5 | 0.00529 | 2.459 | 2.277 |
| NP_000777 | 0.66667 | 3.66667 | 5.5 | 0.0158 | 2.459 | 1.801 |
| NP_002380 | 0.66667 | 3.66667 | 5.5 | 0.0158 | 2.459 | 1.801 |
| NP_001164187 | 0.66667 | 3.66667 | 5.5 | 0.0158 | 2.459 | 1.801 |
| NP_001164187 | 0.66667 | 3.66667 | 5.5 | 0.0158 | 2.459 | 1.801 |
| XP_016869235 | 1.66667 | 9 | 5.4 | 0.01955 | 2.433 | 1.709 |
| NP_005836 | 1 | 5.333333 | 5.333333 | 0.0002 | 2.415 | 3.695 |
| NP_005836 | 1 | 5.333333 | 5.333333 | 0.0002 | 2.415 | 3.695 |
| NP_001185470 | 1 | 5.333333 | 5.333333 | 0.0002 | 2.415 | 3.695 |
| NP_001309896 | 1 | 5.333333 | 5.333333 | 0.00289 | 2.415 | 2.539 |
| NP_001001973 | 1 | 5.333333 | 5.333333 | 0.00289 | 2.415 | 2.539 |
| NP_002296 | 1 | 5.333333 | 5.333333 | 0.00289 | 2.415 | 2.539 |
| NP_714916 | 1 | 5.333333 | 5.333333 | 0.00797 | 2.415 | 2.099 |
| NP_714916 | 1 | 5.333333 | 5.333333 | 0.00797 | 2.415 | 2.099 |
| NP_004544 | 1 | 5.333333 | 5.333333 | 0.00797 | 2.415 | 2.099 |
| NP_000117 | 1 | 5.333333 | 5.333333 | 0.00797 | 2.415 | 2.099 |
| NP_006467 | 1 | 5.333333 | 5.333333 | 0.00797 | 2.415 | 2.099 |
| NP_055755 | 1 | 5.333333 | 5.333333 | 0.02265 | 2.415 | 1.645 |
| NP_036450 | 1 | 5.333333 | 5.333333 | 0.04064 | 2.415 | 1.391 |
| NP_006787 | 1 | 5.333333 | 5.333333 | 0.04064 | 2.415 | 1.391 |
| NP_006787 | 1 | 5.333333 | 5.333333 | 0.04064 | 2.415 | 1.391 |
| NP_001098707 | 1.66667 | 8.666667 | 5.2 | 0.00495 | 2.379 | 2.305 |
| NP_001098707 | 1.66667 | 8.666667 | 5.2 | 0.00495 | 2.379 | 2.305 |
| NP_036313 | 1.33333 | 6.666667 | 5 | 0.0085 | 2.322 | 2.071 |
| NP_001317170 | 0.66667 | 3.333333 | 5 | 0.02322 | 2.322 | 1.634 |
| NP_001616 | 0.66667 | 3.333333 | 5 | 0.04742 | 2.322 | 1.324 |
| NP_001668 | 2 | 10 | 5 | 0.00228 | 2.322 | 2.642 |
| NP_056167 | 1 | 5 | 5 | 0.00228 | 2.322 | 2.642 |
| NP_001014446 | 1 | 5 | 5 | 0.00228 | 2.322 | 2.642 |
| NP_001166925 | 1 | 5 | 5 | 0.01613 | 2.322 | 1.792 |
| NP_001166925 | 1 | 5 | 5 | 0.01613 | 2.322 | 1.792 |
| NP_009034 | 1 | 5 | 5 | 0.01613 | 2.322 | 1.792 |
| NP_057218 | 1 | 5 | 5 | 0.01613 | 2.322 | 1.792 |
| NP_071358 | 2.33333 | 11.33333 | 4.857143 | 0.00882 | 2.28 | 2.054 |
| NP_001139433 | 5.66667 | 27.33333 | 4.823529 | 9.5E-05 | 2.27 | 4.024 |
| NP_006507 | 3.66667 | 17.33333 | 4.727273 | 0.00039 | 2.241 | 3.404 |
| NP_000137 | 14.3333 | 67.66667 | 4.72093 | 0.04029 | 2.239 | 1.395 |
| NP_000144 | 4.33333 | 20.33333 | 4.692308 | 0.00304 | 2.23 | 2.518 |
| NP_003181 | 1 | 4.666667 | 4.666667 | 0.00039 | 2.222 | 3.411 |
| NP_056437 | 1 | 4.666667 | 4.666667 | 0.00039 | 2.222 | 3.411 |
| NP_002309 | 1 | 4.666667 | 4.666667 | 0.00039 | 2.222 | 3.411 |
| NP_689632 | 1 | 4.666667 | 4.666667 | 0.00039 | 2.222 | 3.411 |
| NP_689632 | 1 | 4.666667 | 4.666667 | 0.00039 | 2.222 | 3.411 |
| NP_006347 | 1 | 4.666667 | 4.666667 | 0.038 | 2.222 | 1.42 |
| NP_006347 | 1 | 4.666667 | 4.666667 | 0.038 | 2.222 | 1.42 |
| NP_004125 | 14 | 64.66667 | 4.619048 | 0.00061 | 2.208 | 3.215 |
| NP_596867 | 18 | 82 | 4.555556 | 1E-05 | 2.188 | 4.996 |
| NP_001291217 | 0.66667 | 3 | 4.5 | 0.00219 | 2.17 | 2.659 |
| NP_057086 | 0.66667 | 3 | 4.5 | 0.00219 | 2.17 | 2.659 |
| NP_003835 | 0.66667 | 3 | 4.5 | 0.00219 | 2.17 | 2.659 |
| NP_001309997 | 0.66667 | 3 | 4.5 | 0.0249 | 2.17 | 1.604 |
| XP_011511426 | 1.66667 | 7.333333 | 4.4 | 0.01908 | 2.138 | 1.719 |
| XP_011511426 | 1.66667 | 7.333333 | 4.4 | 0.01908 | 2.138 | 1.719 |
| NP_000427 | 1 | 4.333333 | 4.333333 | 0.00056 | 2.115 | 3.25 |
| NP_000427 | 1 | 4.333333 | 4.333333 | 0.00056 | 2.115 | 3.25 |
| XP_016871724 | 1 | 4.333333 | 4.333333 | 0.00056 | 2.115 | 3.25 |
| XP_016871716 | 1 | 4.333333 | 4.333333 | 0.00056 | 2.115 | 3.25 |
| NP_004246 | 1 | 4.333333 | 4.333333 | 0.00056 | 2.115 | 3.25 |
| NP_056417 | 1 | 4.333333 | 4.333333 | 0.00749 | 2.115 | 2.125 |
| NP_003650 | 1 | 4.333333 | 4.333333 | 0.00749 | 2.115 | 2.125 |
| NP_006342 | 1 | 4.333333 | 4.333333 | 0.00749 | 2.115 | 2.125 |
| NP_056138 | 1 | 4.333333 | 4.333333 | 0.01944 | 2.115 | 1.711 |
| NP_065724 | 2.33333 | 10 | 4.285714 | 0.00802 | 2.1 | 2.096 |
| NP_002071 | 8 | 34 | 4.25 | 0.00105 | 2.087 | 2.981 |
| NP_002071 | 8 | 34 | 4.25 | 0.00105 | 2.087 | 2.981 |
| NP_003312 | 8.33333 | 34.66667 | 4.16 | 0.00015 | 2.057 | 3.817 |
| NP_003312 | 8.33333 | 34.66667 | 4.16 | 0.00015 | 2.057 | 3.817 |
| NP_001070952 | 1.66667 | 6.66667 | 4 | 0.00607 | 2 | 2.217 |
| NP_001001563 | 1 | 4 | 4 | 0.00653 | 2 | 2.185 |
| NP_001254792 | 1 | 4 | 4 | 0.00653 | 2 | 2.185 |
| XP_006715894 | 1 | 4 | 4 | 0.00653 | 2 | 2.185 |
| XP_006715894 | 1 | 4 | 4 | 0.00653 | 2 | 2.185 |
| NP_001317504 | 1 | 4 | 4 | 0.00653 | 2 | 2.185 |
| NP_002990 | 1 | 4 | 4 | 0.00653 | 2 | 2.185 |
| NP_001271342 | 1 | 4 | 4 | 0.00653 | 2 | 2.185 |
| NP_006741 | 1 | 4 | 4 | 0.00653 | 2 | 2.185 |
| NP_006741 | 1 | 4 | 4 | 0.00653 | 2 | 2.185 |
| NP_002817 | 1 | 4 | 4 | 0.00653 | 2 | 2.185 |
| NP_963906 | 1 | 4 | 4 | 0.03994 | 2 | 1.399 |
| NP_004039 | 2.33333 | 9.333333 | 4 | 0.0485 | 2 | 1.314 |
| NP_001327 | 22.3333 | 88.66667 | 3.970149 | 0.00215 | 1.989 | 2.667 |
| NP_001193769 | 13.3333 | 52.66667 | 3.95 | 0.00219 | 1.982 | 2.66 |
| NP_001080 | 10.3333 | 40.66667 | 3.935484 | 0.00026 | 1.977 | 3.581 |
| XP_011532413 | 3 | 11.66667 | 3.888889 | 0.0002 | 1.959 | 3.695 |
| NP_001193769 | 13.3333 | 51.66667 | 3.875 | 0.00241 | 1.954 | 2.618 |
| NP_001104547 | 5 | 19.33333 | 3.866667 | 0.00579 | 1.951 | 2.237 |
| NP_005262 | 4 | 15.33333 | 3.833333 | 0.00921 | 1.939 | 2.036 |
| NP_001269555 | 6.66667 | 24.66667 | 3.7 | 0.00045 | 1.888 | 3.347 |
| NP_001269555 | 6.66667 | 24.66667 | 3.7 | 0.00045 | 1.888 | 3.347 |
| NP_004578 | 9.33333 | 34.33333 | 3.678571 | 0.00202 | 1.879 | 2.694 |
| NP_004578 | 9.33333 | 34.33333 | 3.678571 | 0.00202 | 1.879 | 2.694 |
| NP_001159434 | 1 | 3.666667 | 3.666667 | 0.00132 | 1.874 | 2.878 |
| XP_011536954 | 1 | 3.666667 | 3.666667 | 0.00132 | 1.874 | 2.878 |
| NP_001139630 | 1 | 3.666667 | 3.666667 | 0.00132 | 1.874 | 2.878 |
| NP_733746 | 2 | 7.333333 | 3.666667 | 0.00718 | 1.874 | 2.144 |
| NP_066972 | 1 | 3.666667 | 3.666667 | 0.01613 | 1.874 | 1.792 |
| NP_036335 | 1 | 3.666667 | 3.666667 | 0.03902 | 1.874 | 1.409 |
| NP_036335 | 1 | 3.666667 | 3.666667 | 0.03902 | 1.874 | 1.409 |
| NP_001273300 | 1 | 3.666667 | 3.666667 | 0.03902 | 1.874 | 1.409 |
| NP_001273300 | 1 | 3.666667 | 3.666667 | 0.03902 | 1.874 | 1.409 |
| NP_002547 | 8.66667 | 31.33333 | 3.615385 | 0.00081 | 1.854 | 3.09 |
| NP_005909 | 10 | 35 | 3.5 | 0.0009 | 1.807 | 3.044 |
| NP_001106818 | 12 | 41.66667 | 3.472222 | 9.5E-06 | 1.796 | 5.023 |
| XP_011541265 | 4.33333 | 15 | 3.461538 | 0.00306 | 1.791 | 2.514 |
| XP_011541265 | 4.33333 | 15 | 3.461538 | 0.00306 | 1.791 | 2.514 |
| NP_008879 | 3.33333 | 11.33333 | 3.4 | 0.00194 | 1.766 | 2.713 |
| NP_001094096 | 3.66667 | 12.33333 | 3.363636 | 0.00078 | 1.75 | 3.109 |
| NP_001015 | 1 | 3.333333 | 3.333333 | 0.00219 | 1.737 | 2.659 |
| NP_002812 | 1 | 3.333333 | 3.333333 | 0.00219 | 1.737 | 2.659 |
| NP_000194 | 1 | 3.333333 | 3.333333 | 0.00219 | 1.737 | 2.659 |
| NP_057055 | 1 | 3.333333 | 3.333333 | 0.00219 | 1.737 | 2.659 |
| NP_001273432 | 2 | 6.666667 | 3.333333 | 0.01145 | 1.737 | 1.941 |
| NP_001273432 | 2 | 6.666667 | 3.333333 | 0.01145 | 1.737 | 1.941 |
| NP_002083 | 1 | 3.333333 | 3.333333 | 0.0249 | 1.737 | 1.604 |
| NP_002793 | 1 | 3.333333 | 3.333333 | 0.0249 | 1.737 | 1.604 |
| NP_002793 | 1 | 3.333333 | 3.333333 | 0.0249 | 1.737 | 1.604 |
| NP_001924 | 1 | 3.333333 | 3.333333 | 0.0249 | 1.737 | 1.604 |
| NP_037460 | 1 | 3.333333 | 3.333333 | 0.0249 | 1.737 | 1.604 |
| NP_037460 | 1 | 3.333333 | 3.333333 | 0.0249 | 1.737 | 1.604 |
| NP_002799 | 4 | 13 | 3.25 | 0.00146 | 1.7 | 2.835 |
| NP_002799 | 4 | 13 | 3.25 | 0.00146 | 1.7 | 2.835 |
| NP_060154 | 3.33333 | 10.33333 | 3.1 | 0.00934 | 1.632 | 2.03 |
| NP_060154 | 3.33333 | 10.33333 | 3.1 | 0.00934 | 1.632 | 2.03 |
| NP_001120699 | 7 | 21.66667 | 3.095238 | 0.00365 | 1.63 | 2.438 |
| NP_056083 | 12 | 36.33333 | 3.027778 | 0.00374 | 1.598 | 2.427 |
| NP_001018121 | 3.33333 | 10 | 3 | 0.01301 | 1.585 | 1.886 |
| NP_001280202 | 0.66667 | 2 | 3 | 0.01613 | 1.585 | 1.792 |
| NP_001280202 | 0.66667 | 2 | 3 | 0.01613 | 1.585 | 1.792 |
| NP_000981 | 0.66667 | 2 | 3 | 0.01613 | 1.585 | 1.792 |
| XP_016872987 | 0.66667 | 2 | 3 | 0.01613 | 1.585 | 1.792 |
| NP_001139541 | 0.66667 | 2 | 3 | 0.01613 | 1.585 | 1.792 |
| NP_055621 | 0.66667 | 2 | 3 | 0.01613 | 1.585 | 1.792 |
| NP_001265124 | 1 | 3 | 3 | 0.02572 | 1.585 | 1.59 |
| NP_001138471 | 1 | 3 | 3 | 0.02572 | 1.585 | 1.59 |
| NP_001265392 | 1 | 3 | 3 | 0.02572 | 1.585 | 1.59 |
| NP_001265392 | 1 | 3 | 3 | 0.02572 | 1.585 | 1.59 |
| NP_149100 | 1 | 3 | 3 | 0.02572 | 1.585 | 1.59 |
| NP_001352 | 1 | 3 | 3 | 0.02572 | 1.585 | 1.59 |
| NP_001121132 | 1 | 3 | 3 | 0.02572 | 1.585 | 1.59 |
| NP_001143 | 6 | 17.66667 | 2.944444 | 0.00038 | 1.558 | 3.421 |
| NP_076424 | 3 | 8.666667 | 2.888889 | 0.03251 | 1.531 | 1.488 |
| NP_003642 | 5.66667 | 16.33333 | 2.882353 | 0.04742 | 1.527 | 1.324 |
| NP_001192183 | 2.33333 | 6.666667 | 2.857143 | 0.00436 | 1.515 | 2.361 |
| NP_009216 | 2 | 5.666667 | 2.833333 | 0.02539 | 1.503 | 1.595 |
| NP_877420 | 3.33333 | 9.333333 | 2.8 | 0.00559 | 1.485 | 2.253 |
| NP_001171903 | 1.66667 | 4.666667 | 2.8 | 0.03347 | 1.485 | 1.475 |
| NP_001143 | 7.66667 | 21.33333 | 2.782609 | 0.00039 | 1.476 | 3.404 |
| NP_054735 | 13.6667 | 38 | 2.780488 | 5.3E-05 | 1.475 | 4.276 |
| NP_054735 | 13.6667 | 38 | 2.780488 | 5.3E-05 | 1.475 | 4.276 |
| NP_001627 | 6 | 16.66667 | 2.777778 | 0.00054 | 1.474 | 3.27 |
| NP_001367 | 3 | 8.333333 | 2.777778 | 0.00718 | 1.474 | 2.144 |
| NP_955472 | 53.3333 | 147.3333 | 2.7625 | 0.00247 | 1.466 | 2.607 |
| NP_001189414 | 12 | 33 | 2.75 | 0.00301 | 1.459 | 2.522 |
| NP_113651 | 1.33333 | 3.666667 | 2.75 | 0.03517 | 1.459 | 1.454 |
| NP_003944 | 1.33333 | 3.666667 | 2.75 | 0.03517 | 1.459 | 1.454 |
| NP_001825 | 2 | 5.333333 | 2.666667 | 0.00056 | 1.415 | 3.25 |
| NP_689601 | 1 | 2.666667 | 2.666667 | 0.00749 | 1.415 | 2.125 |
| NP_689450 | 1 | 2.666667 | 2.666667 | 0.00749 | 1.415 | 2.125 |
| NP_001316143 | 1 | 2.666667 | 2.666667 | 0.00749 | 1.415 | 2.125 |
| NP_057292 | 1 | 2.666667 | 2.666667 | 0.00749 | 1.415 | 2.125 |
| NP_001000 | 1 | 2.666667 | 2.666667 | 0.00749 | 1.415 | 2.125 |
| NP_079172 | 1 | 2.666667 | 2.666667 | 0.00749 | 1.415 | 2.125 |
| NP_001245366 | 1 | 2.666667 | 2.666667 | 0.00749 | 1.415 | 2.125 |
| NP_006779 | 3.66667 | 9.666667 | 2.636364 | 0.00129 | 1.399 | 2.888 |
| NP_003036 | 6.33333 | 16.66667 | 2.631579 | 2.6E-05 | 1.396 | 4.591 |
| NP_000099 | 4.33333 | 11.33333 | 2.615385 | 0.00012 | 1.387 | 3.922 |
| NP_789782 | 5 | 13 | 2.6 | 0.04179 | 1.379 | 1.379 |
| NP_001351 | 1.66667 | 4.333333 | 2.6 | 0.04742 | 1.379 | 1.324 |
| NP_710154 | 3 | 7.666667 | 2.555556 | 0.0249 | 1.354 | 1.604 |
| NP_002818 | 3 | 7.666667 | 2.555556 | 0.04881 | 1.354 | 1.311 |
| NP_631918 | 9 | 22.66667 | 2.518519 | 0.00424 | 1.333 | 2.373 |
| NP_001138303 | 5.33333 | 13.33333 | 2.5 | 0.00582 | 1.322 | 2.235 |
| NP_003477 | 3.33333 | 8.333333 | 2.5 | 0.022 | 1.322 | 1.658 |
| NP_006422 | 13.6667 | 34 | 2.487805 | 0.02036 | 1.315 | 1.691 |
| NP_002625 | 7.66667 | 19 | 2.478261 | 0.00105 | 1.309 | 2.979 |
| NP_002625 | 7.66667 | 19 | 2.478261 | 0.00105 | 1.309 | 2.979 |
| XP_016873370 | 7 | 17.33333 | 2.47619 | 0.00207 | 1.308 | 2.685 |
| XP_016873370 | 7 | 17.33333 | 2.47619 | 0.00207 | 1.308 | 2.685 |
| NP_848927 | 3.66667 | 9 | 2.454545 | 0.00377 | 1.295 | 2.423 |
| NP_003833 | 2.33333 | 5.666667 | 2.428571 | 0.02411 | 1.28 | 1.618 |
| NP_001315615 | 25 | 60.66667 | 2.426667 | 0.00238 | 1.279 | 2.624 |
| XP_011534504 | 5 | 12 | 2.4 | 0.00374 | 1.263 | 2.427 |
| NP_002148 | 18.6667 | 44.66667 | 2.392857 | 0.00296 | 1.259 | 2.528 |
| NP_000375 | 5.33333 | 12.66667 | 2.375 | 0.00268 | 1.248 | 2.572 |
| NP_002435 | 5.33333 | 12.66667 | 2.375 | 0.04911 | 1.248 | 1.309 |
| NP_001656 | 3.66667 | 8.666667 | 2.363636 | 0.00045 | 1.241 | 3.349 |
| NP_006030 | 3.66667 | 8.666667 | 2.363636 | 0.00257 | 1.241 | 2.59 |
| XP_005259654 | 1 | 2.333333 | 2.333333 | 0.01613 | 1.222 | 1.792 |
| XP_005259654 | 1 | 2.333333 | 2.333333 | 0.01613 | 1.222 | 1.792 |
| NP_001018080 | 1 | 2.333333 | 2.333333 | 0.01613 | 1.222 | 1.792 |
| NP_001128567 | 1 | 2.333333 | 2.333333 | 0.01613 | 1.222 | 1.792 |
| NP_001311211 | 1 | 2.333333 | 2.333333 | 0.01613 | 1.222 | 1.792 |
| NP_001280134 | 1 | 2.333333 | 2.333333 | 0.01613 | 1.222 | 1.792 |
| NP_000683 | 1 | 2.333333 | 2.333333 | 0.01613 | 1.222 | 1.792 |
| XP_016876295 | 1 | 2.333333 | 2.333333 | 0.01613 | 1.222 | 1.792 |
| XP_016876295 | 1 | 2.333333 | 2.333333 | 0.01613 | 1.222 | 1.792 |
| NP_689518 | 1 | 2.333333 | 2.333333 | 0.01613 | 1.222 | 1.792 |
| NP_919415 | 19.6667 | 45.66667 | 2.322034 | 0.00132 | 1.215 | 2.879 |
| NP_002148 | 20.3333 | 46.66667 | 2.295082 | 0.00212 | 1.199 | 2.674 |
| NP_004550 | 8.66667 | 19.33333 | 2.230769 | 0.00103 | 1.158 | 2.989 |
| NP_001020331 | 3.33333 | 7.333333 | 2.2 | 0.00582 | 1.138 | 2.235 |
| NP_001020331 | 3.33333 | 7.333333 | 2.2 | 0.00582 | 1.138 | 2.235 |
| XP_011529429 | 7.66667 | 16.66667 | 2.173913 | 0.0226 | 1.12 | 1.646 |
| NP_001072989 | 15.6667 | 33.66667 | 2.148936 | 0.04361 | 1.104 | 1.36 |
| NP_004550 | 9.33333 | 20 | 2.142857 | 0.00306 | 1.1 | 2.514 |
| NP_001002814 | 5.66667 | 12 | 2.117647 | 0.00386 | 1.082 | 2.413 |
| XP_005258336 | 46.6667 | 98.33333 | 2.107143 | 0.00015 | 1.075 | 3.825 |
| NP_002061 | 22.6667 | 47.66667 | 2.102941 | 0.0005 | 1.072 | 3.3 |
| NP_001336864 | 4.66667 | 9.666667 | 2.071429 | 0.02846 | 1.051 | 1.546 |
| NP_002363 | 5.33333 | 11 | 2.0625 | 0.02227 | 1.044 | 1.652 |
| NP_002363 | 5.33333 | 11 | 2.0625 | 0.02227 | 1.044 | 1.652 |
| NP_938148 | 34 | 70 | 2.058824 | 0.00051 | 1.042 | 3.292 |
| NP_000503 | 10.3333 | 21 | 2.032258 | 0.03594 | 1.023 | 1.444 |
| NP_000503 | 10.3333 | 21 | 2.032258 | 0.03594 | 1.023 | 1.444 |
| NP_002385 | 23 | 46.33333 | 2.014493 | 4E-06 | 1.01 | 5.4 |
| NP_002385 | 23 | 46.33333 | 2.014493 | 4E-06 | 1.01 | 5.4 |
| NP_061849 | 3.33333 | 6.666667 | 2 | 0.00211 | 1 | 2.676 |
| NP_055422 | 2.33333 | 4.666667 | 2 | 0.00776 | 1 | 2.11 |
| XP_016860258 | 3.66667 | 7.333333 | 2 | 0.00793 | 1 | 2.101 |
| NP_004595 | 2.66667 | 5.333333 | 2 | 0.02322 | 1 | 1.634 |
| NP_004595 | 2.66667 | 5.333333 | 2 | 0.02322 | 1 | 1.634 |

**Supplementary Table 2:**
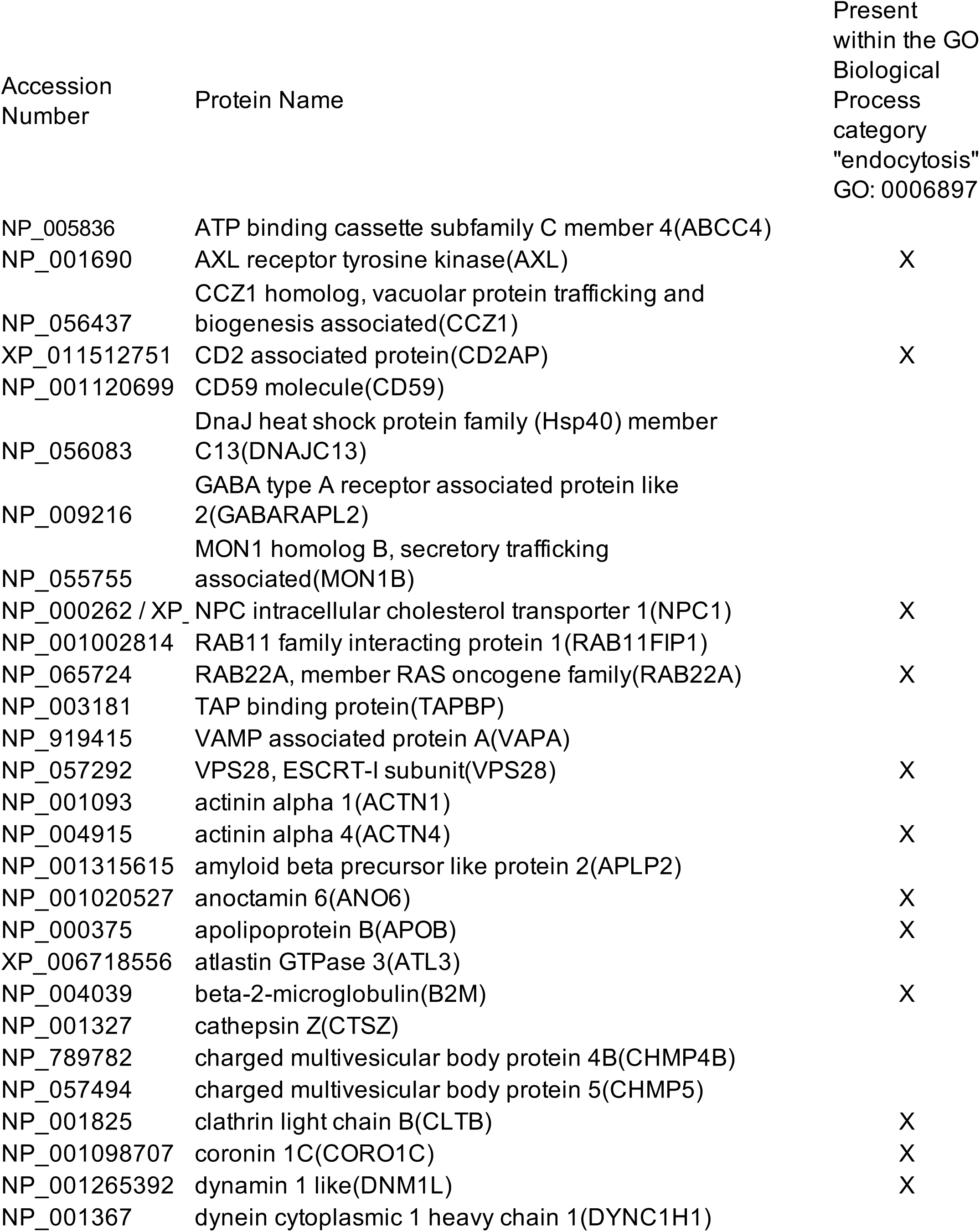
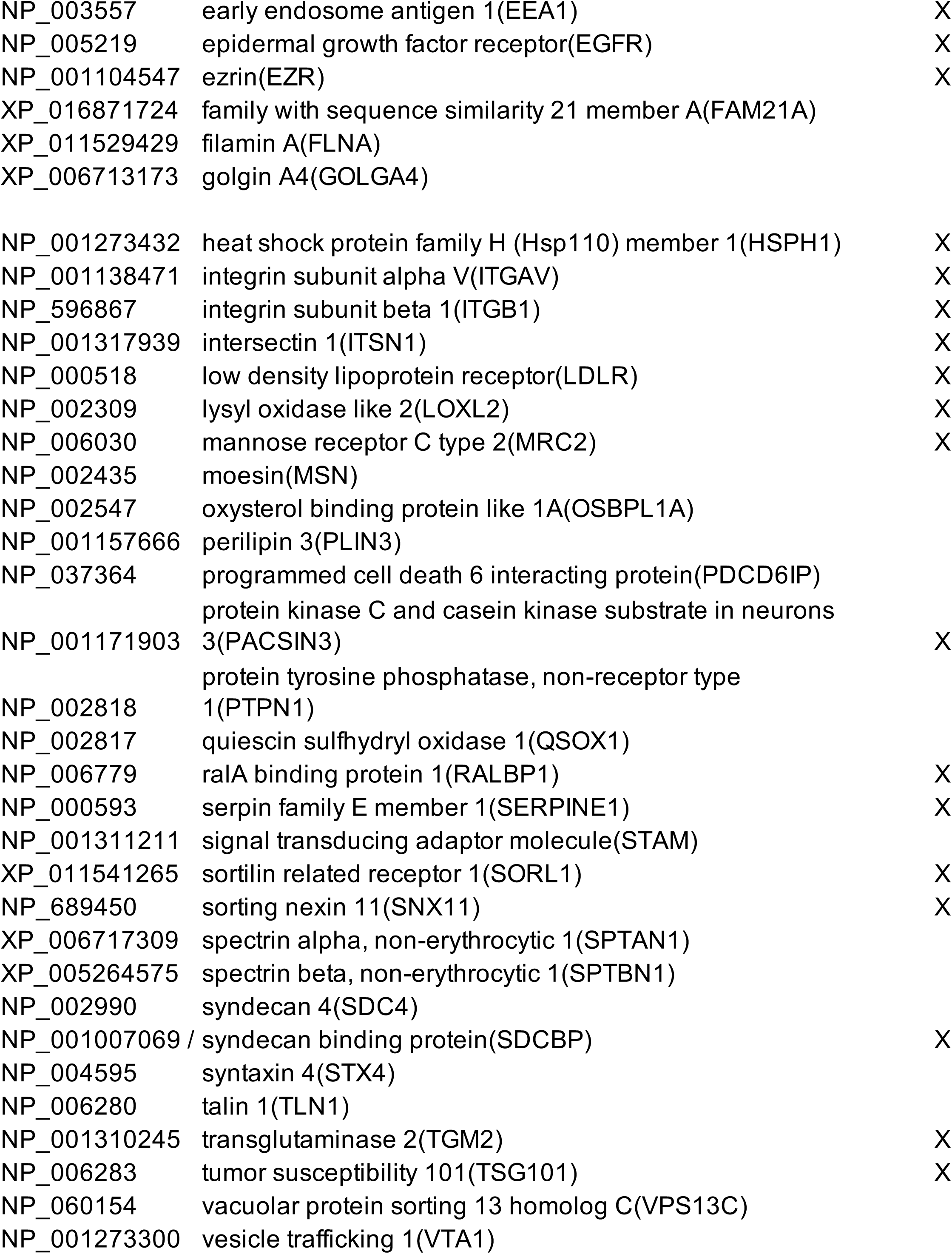
Cohort of *≥*2 fold significantly enriched PDA lysosomal proteins associated with vesicle trafficking and endocytosis.

**Supplementary Table 3:** Clinicopathological characteristics and group membership in ULCA TMA (encompassing resected stage I/II pancreatic cancer)

| Clinicopathologic Category | MYOF IHC staining |  |  |
| --- | --- | --- | --- |
|  | Low | High | p-value<sup>a</sup> |
| **Total patients (n)** | 105 | 31 | N/A |
| **Median Age (yrs)** | 63.7 | 65.4 | N/A |
| **Gender** |  |  |  |
| Male | 63 (60%) | 10 (32%) | 0.006 |
| Female | 42 (40%) | 21 (68%) |  |
| **Histologic grade<sup>c</sup>** |  |  |  |
| Low | 61 (58%) | 14 (45%) | 0.2 |
| High | 44 (42%) | 17 (55%) |  |
| **pT (categorized)** |  |  |  |
| T1+T2 | 17 (16%) | 8 (26%) | 0.29 |
| T2 | 45 (43%) | 9 (29%) |  |
| T3 | 43 (41%) | 14 (45%) |  |
| **pN (categorized)** |  |  |  |
| N0 | 54 (51%) | 12 (39%) | 0.21 |
| N1 | 51 (49%) | 19 (61%) |  |
| **AJCC Stage** |  |  |  |
| I | 33 (31%) | 6 (19%) | 0.19 |
| II | 72 (69%) | 25 (81%) |  |
| **Surgical Margins** |  |  |  |
| R0 | 92 (88%) | 28 (90%) | 0.68 |
| R1 | 13 (12%) | 3 (10%) |  |
| **Tumor size (categorized)** |  |  |  |
| < 3 cm | 64 (61%) | 16 (52%) | 0.35 |
| ≥ 3 cm | 41 (39%) | 15 (48%) |  |

<sup>a</sup> p-values for Chi square test; Low staining =bottom three quartiles (histoscore 0-5); high staining=top quartile (histoscore >5-9). Variables based on the 7th edition of the AJCC/UICC TNM staging system for pancreatic cancer.

**Supplementary Table 4:**
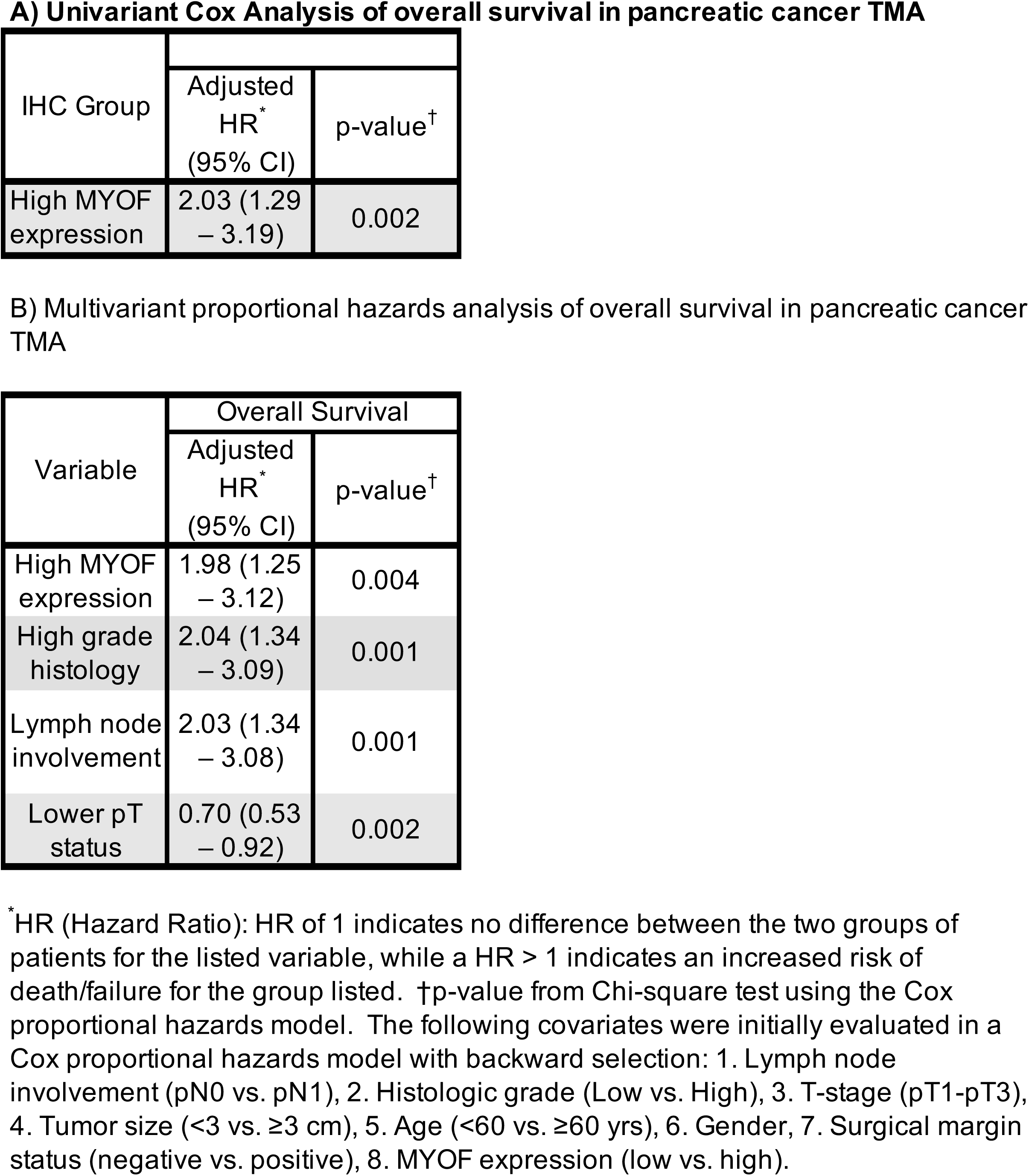
Cox proportional hazard models for prognostic factors.

A) Univariate Cox Analysis of overall survival in pancreatic cancer TMA
| IHC Group |  |  |
| --- | --- | --- |
|  | Adjusted HR*<br>(95% CI) | p-value† |
| High MYOF expression | 2.03 (1.29 – 3.19) | 0.002 |

B) Multivariate proportional hazards analysis of overall survival in pancreatic cancer TMA
| Variable | Overall Survival |  |
| --- | --- | --- |
|  | Adjusted HR*<br>(95% CI) | p-value† |
| High MYOF expression | 1.98 (1.25 – 3.12) | 0.004 |
| High grade histology | 2.04 (1.34 – 3.09) | 0.001 |
| Lymph node involvement | 2.03 (1.34 – 3.08) | 0.001 |
| Lower pT status | 0.70 (0.53 – 0.92) | 0.002 |

\*HR (Hazard Ratio): HR of 1 indicates no difference between the two groups of patients for the listed variable, while a HR > 1 indicates an increased risk of death/failure for the group listed. †p-value from Chi-square test using the Cox proportional hazards model. The following covariates were initially evaluated in a Cox proportional hazards model with backward selection: 1. Lymph node involvement (pN0 vs. pN1), 2. Histologic grade (Low vs. High), 3. T-stage (pT1-pT3), 4. Tumor size (<3 vs. ≥3 cm), 5. Age (<60 vs. ≥60 yrs), 6. Gender, 7. Surgical margin status (negative vs. positive), 8. MYOF expression (low vs. high).

---

**Supplementary Figure 1.**
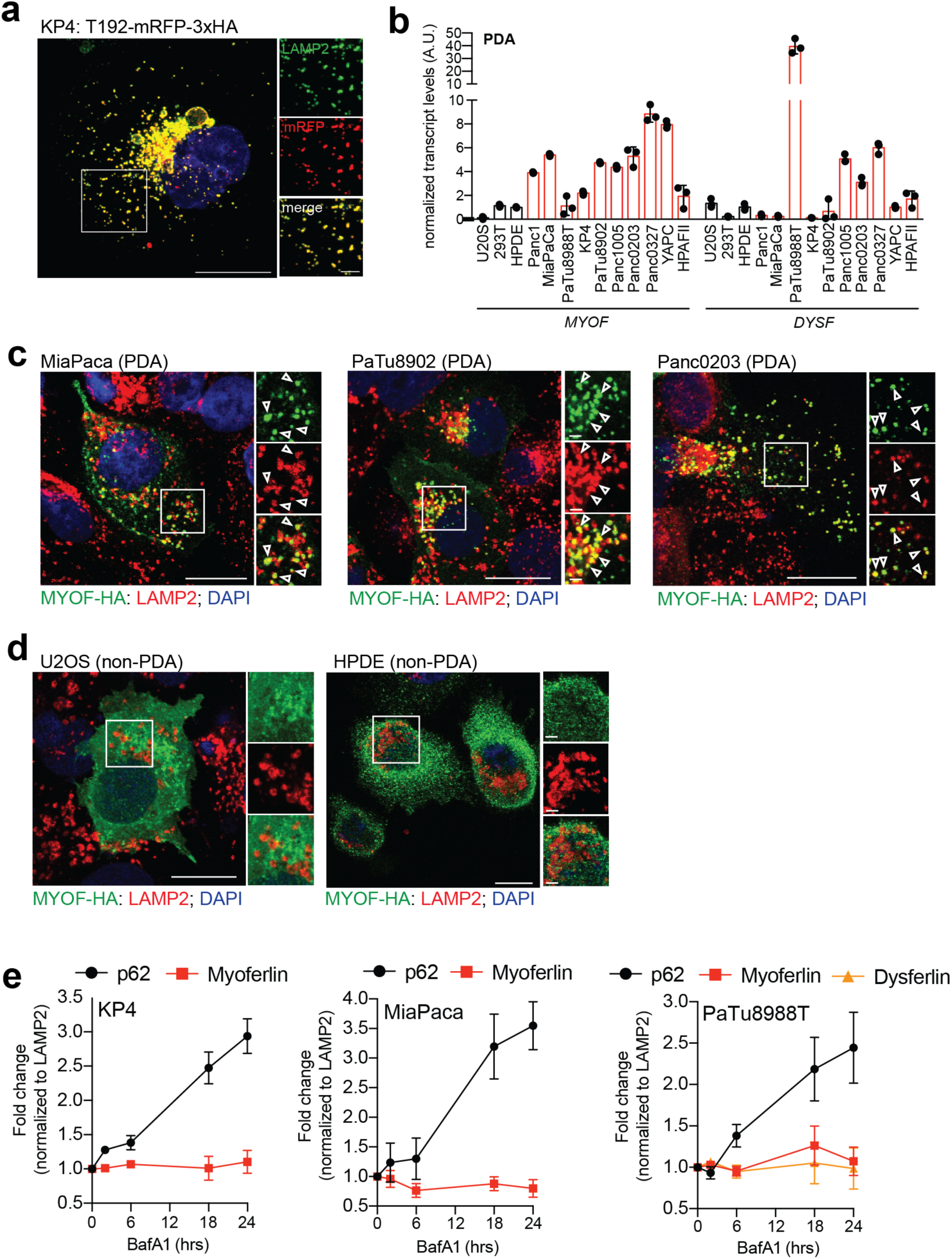
MYOF is a novel lysosomal membrane protein in PDA cells. **a.** Co-localization of T192-mRFP-3xHA and endogenous LAMP2 in KP4 cells. **b.** qRT-PCR analysis of *MYOF* and *DYSF* mRNA levels across 10 human PDA cell lines. **c,d.** Immuno-fluorescence staining of MYOF-HA (green) and LAMP2 (red) in PDA cell lines (MiaPaca, Patu8902 and Panc0203) (c) and non-PDA (U2OS and HPDE) cell lines (d). Arrowheads show examples of co-localization. **e.** Treatment of KP4, MiaPaCa and PaTu8988T cells with 75nM BafA1 for the indicated times causes an increase in p62 levels but not MYOF or DYSF. Graph shows the quantification of normalized fold change relative to LAMP1, averaged from 3 independent experiments. Scale, 20μm for all panels.

**Supplementary Figure 2.**
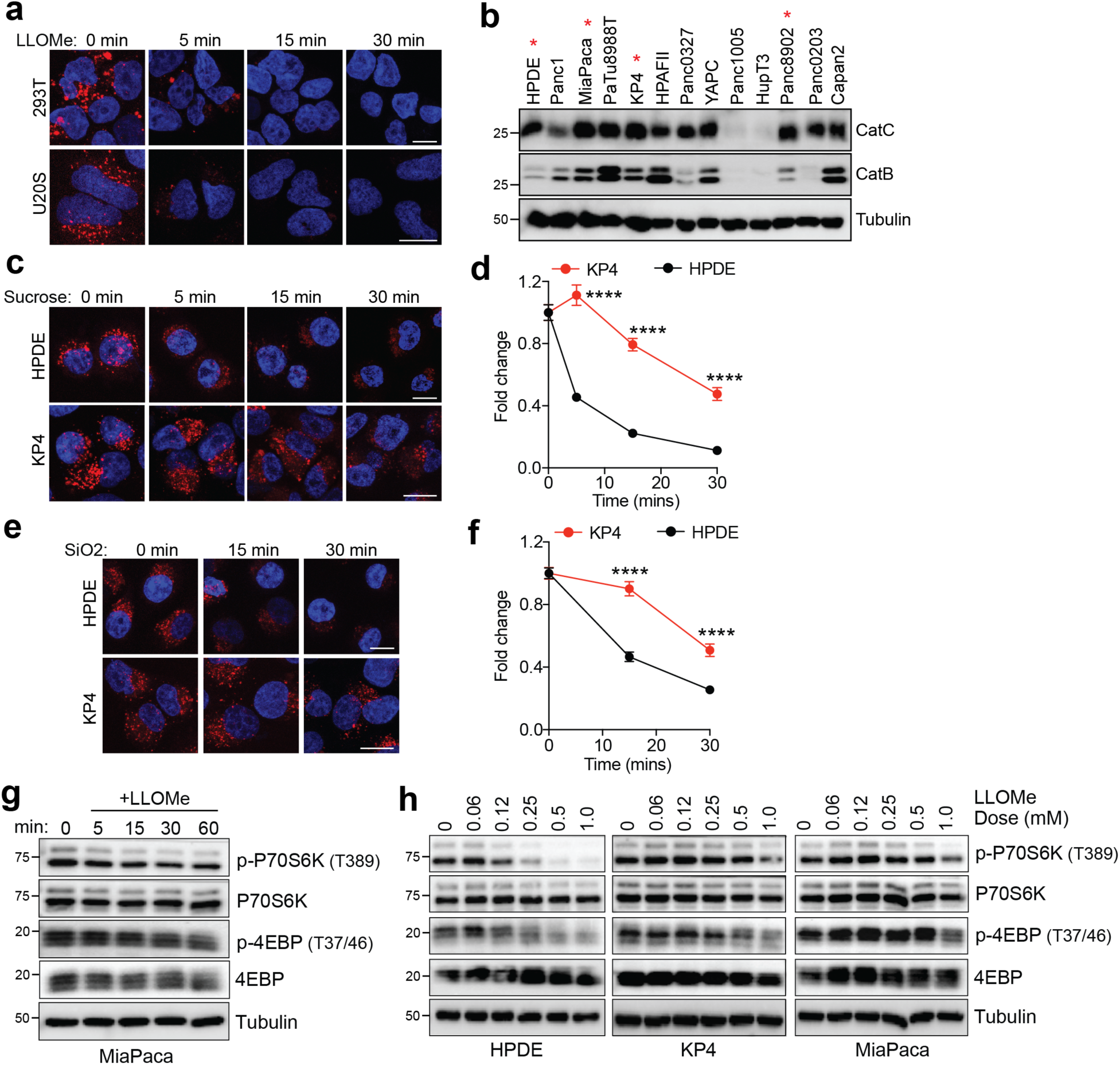
PDA lysosomes are resistant to multiple membrane perturbing agents. **a.** Time-course of lysotracker red staining in 293T and U20S cells following treatment with LLOMe. (293T, n = 80 cells per time point; U20S, n = 57 – 63 cells per time point). **b.** Immunoblot showing the expression of Cathepsin C and Cathepsin B in the indicated cell lines. Asterisk denotes cell lines used throughout the study. **c**. Time-course of lysotracker red staining in HPDE and KP4 cells following treatment with 0.5M sucrose. (HPDE, n = 64 cells per time point; KP4, n = 63 cells per time point). **d.** Normalized fold change of lysotracker staining. **e**. Time-course of lysotracker red staining in HPDE and KP4 cells following treatment with 100 μg/ml silica. (HPDE, n = 65 cells per time point; KP4, n = 65 cells per time point). **f.** Normalized fold change of lysotracker staining. **g.** Immunoblots for the indicated proteins in MiaPaca cells following a time course of 1mM LLOMe treatment. **h.** Immunoblots for the indicated proteins in HPDE, KP4 and MiaPaca cells following treatment for 1hr with increasing doses of LLOMe. Scale, 20μm for all panels. Data are mean ± s.d. *P* values determined by unpaired two-tailed *t*-tests. **** *P* < 0.0001.

**Supplementary Figure 3.**
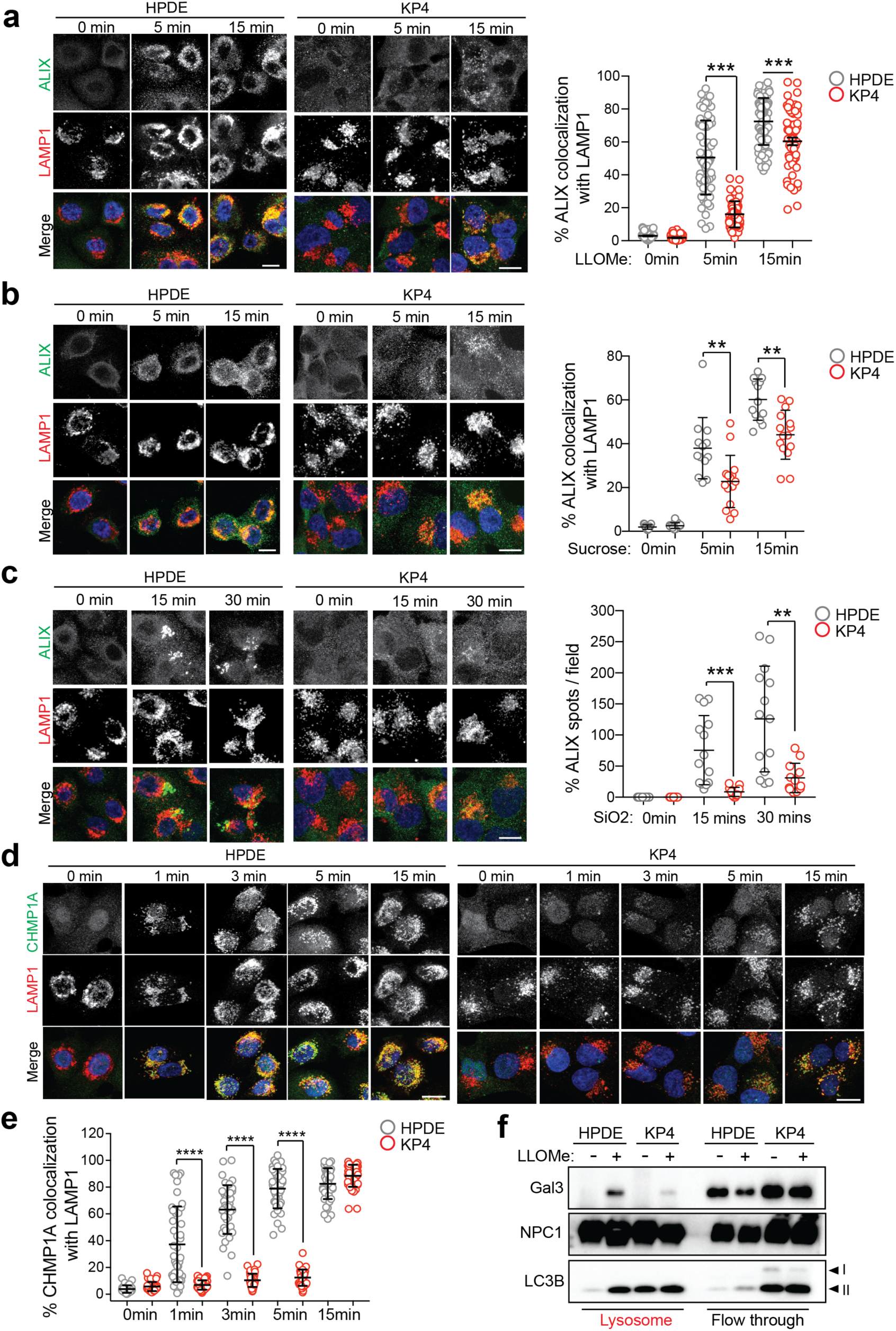
Recruitment of ESCRT proteins to PDA lysosomes is delayed following acute damage. **a-c.** Time course of LLOMe (a), 0.5M sucrose (b), 100 μg/ml silica (c) treatment of HPDE and KP4 cells followed by immuno-fluorescence staining for ALIX (green) and LAMP1 (red). Graphs show the quantification of percentage co-localization of ALIX (LLOMe, n = 60; sucrose, n = 13-15 fields/conditions; silica, n = 12-14 fields/condition per cell line) with LAMP1 positive lysosomes. **d.** Time course of LLOMe treatment of HPDE and KP4 cells followed by immuno-fluorescence staining for CHMP1A (green) and LAMP1 (red). **e**. Graph shows quantification of percentage co-localization of CHMP1A (n = 40 per cell line) with LAMP1 positive lysosomes. **f.** Immunoblot for the indicated proteins in lysosome fractions and flow through fractions isolated from HPDE- and KP4-T192-mRFP-3xHA stable cell lines treated with LLOMe for 10min. Scale, 20μm. Data are mean ± s.d. *P* values determined by unpaired two-tailed *t*-tests. ** *P* < 0.01; *** *P* < 0.001; **** *P* < 0.0001.

**Supplementary Figure 4.**
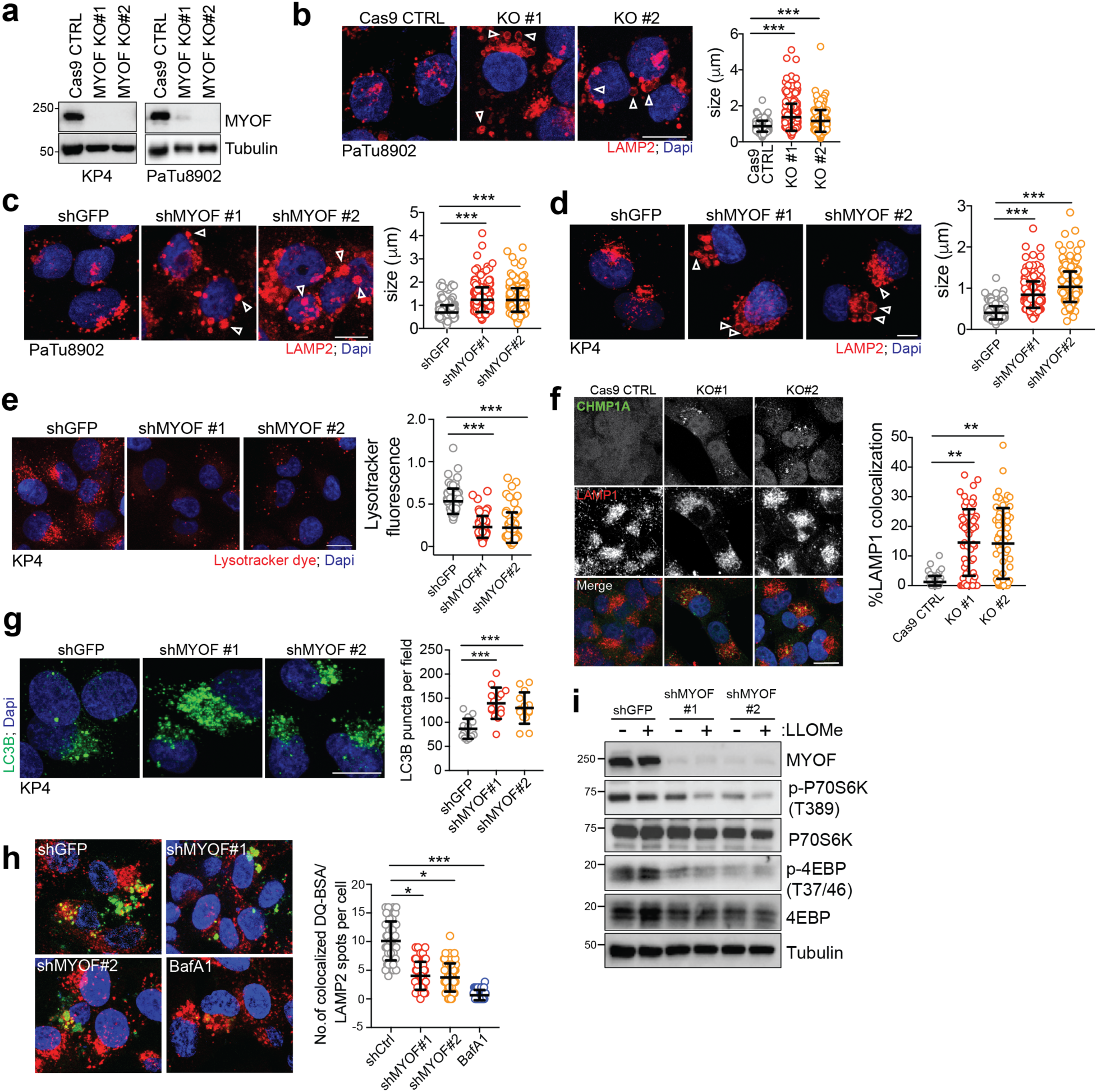
MYOF loss leads to lysosome dysfunction in PDA cells. **a.** Immunoblot showing the efficiency of MYOF ablation in KP4 and PaTu8902 CRISPR KO clones. **b.** MYOF KO in PaTu8902 cells causes aberrant lysosomal morphology and increased size as shown by LAMP2 staining (arrowheads). Graph on the right shows the measurement of lysosome diameter in Cas9 control cells (n = 252), MYOF KO #1 (n = 255) and MYOF KO #2 (n = 258). **c, d.** shRNA mediated knockdown of MYOF in PaTu8902 (c) and KP4 (d) cells leads to defects in lysosome morphology and size as shown by LAMP2 staining (arrowheads). Graphs on the right show the measurement of lysosome diameter [8902 shGFP (n = 258), shMYOF#1 (n = 256), shMYOF#2 (n = 263); KP4 shGFP (n = 254), shMYOF#1 (n = 239), shMYOF#2 (n = 243) cells]. **e.** Lysotracker staining in KP4 cells following infection with shRNA against GFP or MYOF. Graph on the right shows the quantification of normalized fold change in lysotracker fluorescence in control (shGFP, n = 70 cells) and KD (shMYOF#1, n = 65; shMYOF#2, n = 77) conditions. **f.** Increased recruitment of CHMP1A (green) to LAMP1 positive lysosomes (red) following KO of MYOF (#1, n = 60; #2, n = 61) in KP4 cells relative to Cas9 control (n = 58) cells. Graph shows the quantification of percentage CHMP1A co-localization with LAMP1. **g.** Immuno-fluorescence staining for LC3B in PaTu8902 cells following infection with shGFP (n = 55 cells) or shRNA mediated knockdown of MYOF (#1, n = 57; #2, n = 58 cells). Graph on the right shows the quantification of LC3B puncta. **h.** Proteolysis of macropinocytosed protein is impaired following shRNA mediated knockdown of MYOF or BafA1 treatment as determined by pulse-chase with DQ-BSA. Degradation of DQ-BSA in lysosomes is quantified (number of fluorescent spots/cell co-localizing with LAMP2 positive lysosomes from n = 54 (shGFP), 57 (shMYOF#1), 52 (shMYOF#2), 54 (BafA1 treated) cells. **i**. Immunoblots for the indicated proteins in KP4 control cells (shGFP) or following shRNA mediated knockdown of MYOF, with or without 1mM LLOMe treatment to determine mTORC1 signaling activity. Data are mean ± s.d. Scale, 20μm for all panels. *P* values determined by unpaired two-tailed *t*-tests. * *P* < 0.05; ** *P* < 0.01; *** *P* < 0.001; **** *P* < 0.0001.

**Supplementary Figure 5.**
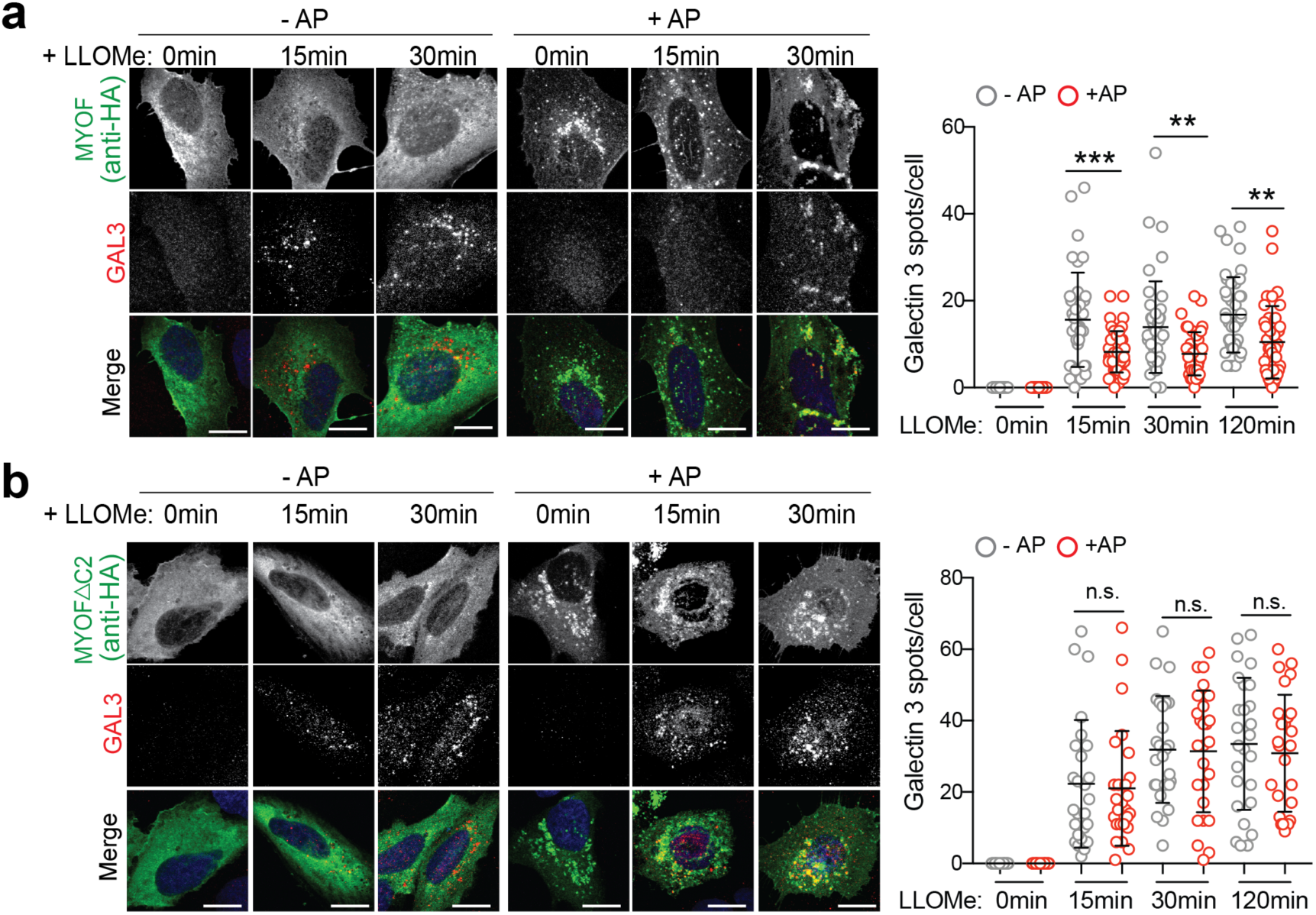
Lysosomal targeting of MYOF delays onset of membrane damage. **a.** U20S cells stably expressing T192-Flag-FKBP and transiently transfected with MYOF-FRB* were treated with 1mM LLOMe for the indicated time points in the presence or absence of AP, followed by immuno-staining for HA (green) and Galectin 3 (GAL3; red). Recruitment of MYOF-FRB* (green) protects against LLOMe induced Gal3 recruitment [n= 40 (control), 39 (15mins), 43 (30 mins), 41 (120mins) cells in the absence of AP and n = 39 (control), 45 (15mins), 41 (30mins) and 39 (120mins) cells in the presence of AP]. **b.** U20S cells stably expressing T192-Flag-FKBP and transfected with MYOF*Δ*C2-FRB* variant were treated as in ‘a’, followed by immuno-staining for HA (green) and ALIX (red). Recruitment of MYOF*Δ*C2 (green) does not protect against LLOMe induced ALIX recruitment [n = 26 cells per condition (-AP and +AP)]. Graphs at right show quantification of GAL3 (top) and ALIX (bottom) spots per cell in response to LLOMe. Scale, 20μm for all panels. Data are mean ± s.d. *P* values determined by unpaired two-tailed *t*-tests. ** *P* < 0.01; *** *P* < 0.001; n.s. not significant.

**Supplementary Figure 6.**
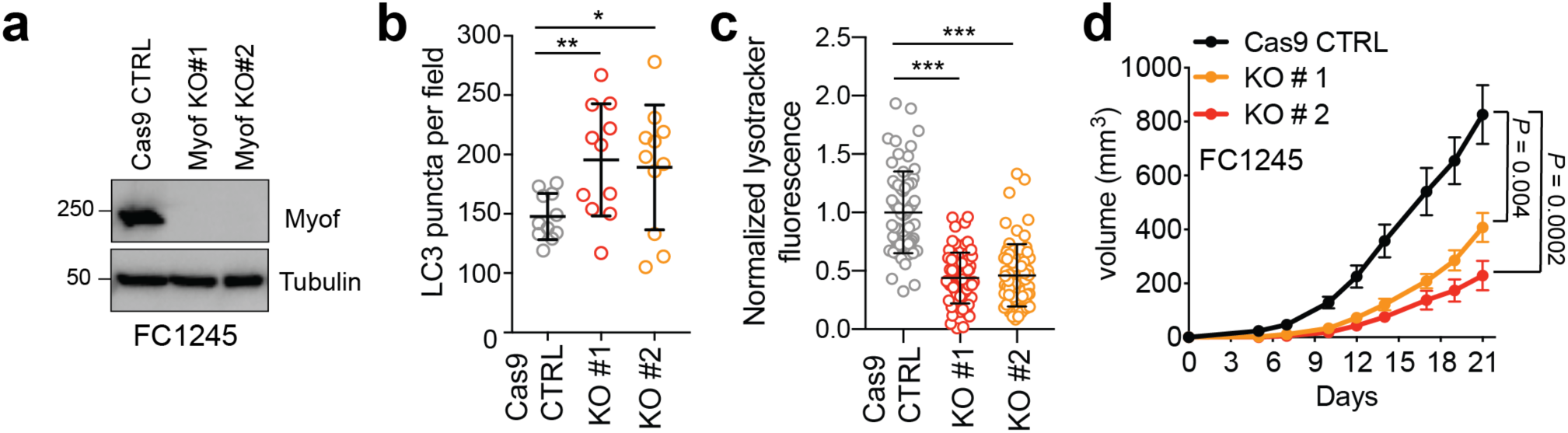
MYOF is required for PDA tumour growth. **a.** Immunoblot of the indicated proteins in mouse KPC cells (FC1245) following CRISPR mediated knockout of Myof. **b**. Graph showing increased accumulation of LC3B positive autophagosomes in FC1245 Myof KO cells quantified from n = 11 fields per condition. Data are mean ± s.d. **c.** Graph showing decrease in lysotracker red staining in FC1245 Myof KO cells compared to Cas9 control cells, quantified from n = 70 (Cas9 control), 71 (KO#1), 70 (KO#2) cells per condition. Data are mean ± s.d. **d.** Growth rate of Cas9 control and Myof KO FC1245 allografts following s.c. transplantation in syngeneic C57BL/6 host mice (n = 5-6 animals per group). Data represent mean ± s.e.m. *P* values determined by unpaired two-tailed *t*-tests. * *P* < 0.05; ** *P* < 0.01; *** *P* < 0.001.

